# Genome scan of landrace populations of the self-fertilizing crop species rice, collected across time, revealed climate changes’ selective footprints in the genes network regulating flowering time

**DOI:** 10.1101/2022.07.29.502004

**Authors:** Nour Ahmadi, Mamadou Billo Barry, Julien Frouin, Miguel de Navascués, Mamadou Aminata Toure

## Abstract

Analysis of the genetic bases of adaptation to climate changes are often conducted on natural populations. We report here on a study based on diachronic sampling (1980 & 2010) of the self-fertilising crop species, *Oryza sativa* (Asian rice) and *Oryza glaberrima* (African rice), in the tropical forest and the Sudanian savannah of West Africa. First, using historical meteorological data we confirmed changes in temperatures (+1°C on average) and rainfall regime (less predictable and reduced amount) in the target area. Second, phenotyping the populations for phenology, we observed significantly earlier heading time (up to 10 days) in the 2010 samples. Third, we implemented two genome-scan methods, one of which specially developed for selfing species, and detected 31 independent selection footprints. These loci showed significant enrichment in genes involved in reproductive processes and bore known heading time QTLs and genes, including *OsGI, Hd1* and *OsphyB*. This rapid adaptive evolution, originated from subtle changes in the standing variation in genetic network regulating heading time, did not translate into predominance of multilocus genotypes, as it is often the case in selfing plants, and into notable selective sweeps. We argue that this high adaptive potential results from the multiline genetic structure of the rice landraces, and the rather large and imbricated genetic diversity of the rice meta-population at the farm, the village and the region levels, that hosted the adaptive variants in multiple genetic backgrounds well before the advent of the environmental selective pressure. The complex selection footprints observed in this empirical study calls for further model development on genetic bases of plant adaptation to environmental changes.

## Introduction

Climate change is affecting agricultural productivity and its effects may limit global crop production by at least 10% in 2050 (Davis et al., 2016), especially in farming systems dominated by rainfed crops such as in sub-Saharan Africa (Sultan et al., 2019). Adaptation strategies include using better-adapted varieties or switching to more adapted crop species (Pironon et al., 2019). The variety replacement strategy relies on selection of better-adapted varieties among the ones currently cultivated or maintained in gene banks (Rojas, 2019), and on creation of new varieties combining adaptive traits scattered among the cultivated species (Haussmann et al., 2012; Watson 2019) and their wild relatives (Cortés & López-Hernández, 2021).

Climate change also imposes strong selective pressure on natural populations and on standing genetic diversity of crop species. Predicted consequences include species range shifts, altered phenology, and local extinction (Reush & Wood, 2007). Understanding how climate changes affect phenotypic and genetic diversity of crop species can provide clues on which type of material we should produce to withstand the new climatic regimes (Snowdon et al., 2021). It can also provide information on the species vulnerability to these environmental changes (Hoffmann & Willi, 2008).

Three mechanisms are commonly distinguished as possible responses to climate change: migration, phenotypic plasticity and genetic adaptation (Hoffmann & Sgrò, 2011; Franks et al., 2014). It has been shown that for many plant species the migration pace cannot follow the pace of predicted local climate change (Lorie et al., 2009). Adaptation can occur on a relatively short time if the plant population has sufficient genetic variation upon which selection can act (Hoffmann & Sgrò, 2011; Pauls et al., 2013; Messer & Petrov, 2013). Organisms can also respond to climate change through phenotypic plasticity, i.e., the ability of a given genotype to modify its phenotype in response to environmental variation (Springate et al., 2013; Kelly, 2019). A major unknown is the relative importance of genetic adaptation versus phenotypic plasticity in responding to rapid environmental change and the way they interact (Messer & Petrov, 2013).

Understanding mechanisms involved in genetic adaptation linked to spatial or temporal heterogeneity in selection pressures is also an important issue of evolutionary and conservation biology. This implies knowledge about the role of evolutionary processes such as mutation and recombination (Reush & Wood, 2007), and about the relative importance of standing genetic variation (i.e. allelic variation that is currently segregating within a population or a species) as opposed to new mutations, as a source of beneficial alleles (Hermisson & Pennings, 2017). The role of selection in the adaptation process is also critical to understand, as it shapes the evolution of genes involved in adaptive traits as well as their surrounding genetic diversity (reviewed in Orr, 2002; 2005). Answers to these questions may also depend on the biological characteristics of the species under consideration (life cycle, mating system, dispersal rate, etc.). For instance, adaptation to new conditions is expected to be limited in predominantly self-fertilizing populations, due to reduced effective population size and recombination (Anderson et al., 2011; Hartfield et al., 2017). It is thus important to get empirical data on adaptation routes and genetic mechanisms in both self-fertilizing and outcrossing species. Geographic areas where farming systems, not yet affected by the Green Revolution, still rely on a large number of landraces constitute a good setting for such empirical analysis of the genetics of crop species adaptation to climate changes. Indeed, when spread over a broad environmental gradient, those landraces can collectively represent a meta-population harbouring large standing genetic variation (Barry et al., 2007c; Radanielina et al., 2013).

Plants phenotypic response to climate changes is as multi-faceted as the changes in environmental parameters. Among the plant phenotypic traits, the reproductive development process and phenology are particularly responsive to climate changes (Gray & Brady, 2016; Prevéy, 2020). The initiation of the reproduction phase is a critical life history step and its timing has important fitness consequences. Flowering too early or too late can induce sterility and reduced fecundation rate due to harsh conditions such as high temperatures (Lohani et al., 2020), or incomplete seed development due to adverse meteorological phenomenons such as drought (Fahad et al., 2017; Yu et al., 2019). Regulation of flowering time is a complex process often involving several genetic pathways. In *Arabidopsis thaliana*, it involves a network of more than 60 genes, regulated through four different pathways (Wellmer & Riechmann, 2010; Quiroz et al., 2021). In rice (*Oryza sativa*) the list of known regulators of flowering time (termed heading date) includes some 53 genes involved in at least three pathways: photoperiod and circadian clock, chromatin-related pathway, and hormones pathway (Wei et al., 2020).

Several approaches have been developed for the analysis of the genetic bases of adaptation to climate changes (reviewed in Franks & Hoffmann, 2012; Aguirre-Liguori et al., 2021) including artificial evolutionary experiments (Hansen et al., 2012), sampling along cline (Bailey & Bataillon, 2016; Cruzan & Hendrickson, 2020), or sampling across time (Franks & Weis, 2008; Hansen et al., 2012) associated with genome scans techniques. However, almost all of the latter studies were focused on natural populations (Bailey & Bataillon, 2016).

Here we present an analysis of the genetic bases of adaptation to climate changes based on sampling across time, in an autogamous crop species, rice, of major significance for the world food security. Our study place was the tropical forest and the Sudanian savannah areas of Guinea in West Africa. The interval between the two sampling times was approximately 30 years. First, using local historical meteorological data we analysed and confirmed the reality of climate changes in the target area. Second, we analysed and confirmed significant phenological differences between the populations of the Asian rice (*Oryza sativa*) and of the African rice (*Oryza glaberrima*) sampled across time. Third, we implemented two genome-scan methods on those populations and detected several selection footprints. Finally, we were able to connect the loci under selection to known genes and QTLs involved in reproductive development. Our results provide new insight into mechanisms of adaptation of self-fertilizing plant meta-populations, under environmental and human selection.

## Materiel and methods

### Study periods, geographic area, and collection strategy

Plant materiel was collected through two collection campaigns in Conakry Guinea. The first campaign (hereafter referred to as Collect-1) took place in 1979 and 1982 (Bezançon et al., 1983). The second collection campaign (Collect-2) was undertaken in 2011 by scientists from Cirad (https://www.cirad.fr/) and IRAG (https://irag-guinee.org/). The collect area stretched from tropical forest to Sudanian savannah agro-ecological regions of Conakry Guinea: latitude 7°34’19.96”N-12°29’1.50”N; longitude 8°25’0.0”O-13°17’59.17”O (S-Figure 1).

The collect strategy was to drive along major roads and other practicable trails, stop in villages without prior notice, conduct a quick inventory of the rice varieties and collect a sample of each of the identified varieties, in the field or from the granary. The passport data for Collect-1 samples included the village name, the variety name, the species (*O. sativa* or *O. glaberrima*) and, occasionally, the rice growing ecosystem (upland, hydromorphic, lowland) and the type of collect (field/granary, bulk/panicle). Approximately 40% of the accessions were collected in the field.

To prepare the Collect-2 campaign, the geographic coordinates of a large share of the villages visited during Collect-1 was retrieved using Google-Earth software. The collect team borrowed the track of Collect-1 campaign but did not limit its collect to the Collect-1 villages, especially as some villages could not be found. The passport data for samples from Collect-2, included the village name and its geographic coordinates, the variety name and tentative species membership, the rice growing ecosystem and the type of collect. Approximately 80% of the samples were panicles collected in the field.

For Collect-1, 442 accessions were retrieved from the long-term conservation facilities of the Institute de Recherche pour le Development (https://www.ird.fr/). The Collect-2 campaign yielded 776 accessions.

### Genotyping

All Collect-1 and Collect-2 accessions were first genotyped with 16 SSR markers using DNA from a plant obtained with a seed visually most representative of the seed stock of each accession. These data served to (i) ascertain the membership of accessions to one of the three genetic groups (*O. sativa indica, O. sativa japonica*, and *O. glaberrima*), (ii) detect redundant accessions, (iii) assess the pattern of genetic diversity within each genetic group, for each collect. Based on such information and logistic issues, 600 accessions were selected for genotyping by sequencing (GBS). DNA libraries were prepared at the Regional Genotyping Technology Platform (http://www.gptr-lr-genotypage.com), using the MATAB method for DNA extraction and the *ApekI* enzyme for DNA digestion. The libraries were single-end sequenced using Illumina HiSeq™2000 (Illumina, Inc.) at the Regional Genotyping Platform (http://get.genotoul.fr/). The Fastq sequences were aligned to the rice reference genome, Os-Nipponbare-Reference-IRGSP-1.0 with Bowtie2 (default parameters). Non-aligning sequences and sequences with multiple positions were discarded. Single nucleotide polymorphisms (SNP) were called using Tassel GBS pipeline v5.2.29 (Bradbury et al., 2007). First, the genotypic data for all accessions taken together were filtered for quality score (> 20) and bi-allelic status of SNPs. Second, the genotypic data of accessions from each genetic group were filtered separately for minor allele frequency (MAF > 1%), rate of missing data (< 20%), and heterozygosity (< 5%). The missing data were imputed, separately for each group, using Beagle v4.0. Table 1 summarizes the results of the process.

**Table 1:** Rice accessions from the two collect campaigns that were genotyped and/or phenotyped for the present study.

| Genetic group | Number of SNP | Number of accessions genotyped |  |  | Number of accessions phenotype |  |  |
| --- | --- | --- | --- | --- | --- | --- | --- |
|  |  | Collec-1 | Collect-2 | Total | Collect-1 | Collect-2 | Total |
| *O. sativa Indica (Osi)* | 14,775 | 15 | 55 | 70 | 73 | 180 | 253 |
| *O. sativa japonica tropical (Osj)* | 9,282 | 96 | 154 | 250 | 107 | 147 | 254 |
| *O. glaberrima (Og)* | 6,250 | 111 | 99 | 210 | 59 | 85 | 144 |
| Total |  | 222 | 308 | 530 | 239 | 412 | 651 |

### Phenotyping

Independent from the above-mentioned selection of accessions for genotyping, 239 accessions from Collect-1 and 412 accessions from Collect-2 were selected (Table 1) to be phenotyped for the duration of the sowing to heading date (DTHD). Accessions from Collect-1 were first seed increased in a greenhouse in Montpellier, France. Phenotyping of accessions collected from the upland and the lowland ecosystems was performed separately. The 362 accessions collected from the upland were cultivated in an upland field, in the IRAG research station in Guinea (Kindia, 10°00’51.37 N, 12°50’66” W), in 2012. Accessions grown in the lowland were sown twice (mid-July and mid-August) in 2013. The elementary plot for each accession was three consecutive lines of 1m long, and the DTHD was recorded on five plants of the middle line. Homogeneity of growing conditions was assessed by the systematic presence of two check varieties every 20 test accessions. There were 364 accessions in common between GBS genotyping and field phenotyping.

### Assessment of genetic diversity and population parameters

Pattern of genetic diversity at the whole accession level and, separately, at the level of each genetic group was investigated using distances between individual accessions estimated by a simple-matching dissimilarity index, computed from genotypes at 1,130 SNP common to the *O. sativa indica* (*Osi*), *O. sativa* tropical *japonica* (*Osj*) and *O. glaberrima* (*Og*) accessions. An unweighted neighbour-joining tree was constructed based on this dissimilarity matrix, using DARwin.6 (Perrier & Jacquemond-Collet, 2006).

Population parameters were assessed for each genetic group subdivided into two subpopulations representing Collect-1 and Collect-2. This is *Osi*-1 and *Osi*-2 for the *O. sativa indica, Osj*-1 and *Osj*-2 for the *O. sativa* tropical *japonica*, and *Og*-1 and *Og*-2 for *O. glaberrima*. The relative importance of genetic diversity among subpopulations, among individuals within a subpopulation and within individuals, was assessed by analysis of molecular variance AMOVA with Arlequin software version 3.5.2. (Excoffier et al., 2010). The same software served to compute nucleotide diversity, expected heterozygosity per locus, the Wright (1965) F-statistics (*F*_*IS*_ and *F*_*ST*_) and the population parameter *θ* = 4*N*_*e*_μ, where *N*_*e*_ is the effective population size and µ is the overall mutation rate at the haplotype level.

The speed of decay of linkage disequilibrium (LD) in each subpopulation was estimated by computing r^2^ between pairs of markers on a chromosome basis, using the “full matrix” option of Tassel 5.2.6 (Bradbury et al., 2007), and then by averaging the results by classes of distance.

### Detection of loci under selection

Presence of loci under selection was investigated through genome scan for loci for which the extent of differentiation between subpopulations Collect-1 and Collect-2 of a given genetic group, summarized by the *F*_*ST*_ statistic, is significantly higher than the expected *F*_*ST*_ values under neutrality hypothesis, for a given average heterozygosity (*h*_0_) between populations and for a given demographic model (Beaumont & Nichols (1996). Two genome scan methods were implemented: a standard approach assuming outcrossing reproduction (Excoffier et al., 2009) and a method specifically developed for mainly autogamous species (Navascués et al., 2021).

In the Excoffier et al. (2009) approach (hereafter referred to as the “heterozygosity-based” method), first the locus-by-locus differentiation *F*_*STi*_ is computed as 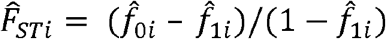, where 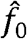 is the average homozygosity within a population and 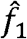 is the probability that two genes from different subpopulations are identical (Weir & Cockerham (1984). Second, Global *F*_*ST*_ is computed as a weighted average among loci, where 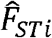 values are weighed by the heterozygosity between populations 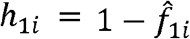 and then the heterozygosity between populations 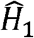 is inferred from the average heterozygosity 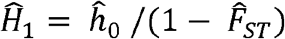. Third, the joint distribution of the neutrality *F*_*ST*_ and heterozygosity is obtained by coalescent simulation under the hypothesis of hierarchical island model of population differentiation. Finally, locus-specific *F*_*ST*_ P-values are obtained from the simulated joint distribution of 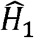 and 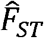 by a kernel density approach. Under the hierarchical island model, the population structure is arranged into *k* groups of *d* demes, in which migration rates between demes are different within and between groups. The mutation model implemented is the SNP mutation model, in which a mutation occurs at random along the simulated structured coalescent tree. The approach was implemented using Arlequin 3.5.2 (Excoffier et al., 2010). The value for *k* and *d* was set to 50 and 10, respectively, to mimic a large number of villages each owning less than 10 rice varieties and preferential interactions between varieties of the given village. The number of simulations was set to 50,000.

The Navascués et al. (2021) method (hereafter referred to as “drift-based” method), relies on principles first proposed by Goldringer & Bataillon (2004), that is using the estimated magnitude of drift between two time samples as a null model to test for homogeneity of differentiation across loci. In the case of complete neutrality, all sampled loci should provide estimates of genetic differentiation drawn from the same distribution. Assuming a single isolated population, this distribution depends on the strength of the genetic drift, that is, on the length of the period (*t* in number of generations) and on the effective population size (*N*_*e*_). Loci under directional selection or linked to such loci are expected to exhibit larger differentiation values than expected under the neutral hypothesis. In order to account for the mainly selfing reproduction system, Navascués et al. (2021) developed an implementation procedure that relies not on the observed allele frequency, but on genotype (*AA, Aa* and *aa*) frequency. Indeed, in a mainly selfing species, allele copies within an individual are not independent samples from the population, while genotypes are independent samples. Moreover, in order to account for uncertainty of the initial genotype frequency, assuming the same prior probability for the three genotype frequencies, genotype frequencies in the population are sampled from the posterior probability distribution with Dir(K_0_ + 1), where K_0_ is the observed genotype counts in the sample of the focal locus at time *t* = 0.

Locus-by-locus differentiation *F*_*STi*_ are computed with the Weir & Cockerham (1984) method. Temporal differentiation is measured by estimating the *F*_*ST*_ with the analysis of variance approach proposed by Weir & Cockerham (1984). Then, the temporal *F*_*ST*_ is used to estimate *N*_*e*_ as: 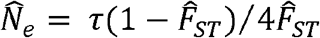, where *τ* is the number of generations between the two temporal populations. The null distribution of single locus *F*_*STi*_ is built through simulations of drift. Each of these simulations consists in (i) drawing the initial genotype frequency K_0_ of the locus, conditional on data, (ii) simulating allele frequency change for *τ* generations, based on 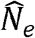, (iii) simulating samples by sampling genotypes with genotype frequencies based on 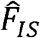, and (iv) calculating 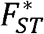 for the simulated sample. The proportion of 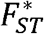 equal or larger than the observed 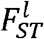 provides an estimate of the *p*-value for the test. In order to account for multiple tests across the genome, false discovery rates (q-values) are computed from the distribution of the p-value. Genotype counts observed in the simulated sample of n_0_ individuals are modelled as coming from a multinomial distribution Mult(n_0_, *γ*_0_), where *γ*_0_ are the genotype frequencies in the population. In the subsequent generations, allele frequencies *π*_*t*_ are simulated following a binomial distribution as 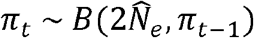 where 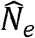 is the genome wide estimate of the effective population size and *π*_0_ is determined by *γ*_0_. At time *t* = τ, simulated genotype counts in samples are taken from the multidimensional distribution 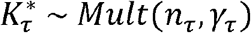, were *n*_*τ*_ is the size of the sampled population at that time, and *γ*_*τ*_ are the genotype (*AA, Aa* and *aa*) frequencies in the populations as a function of the allele frequency *π*_*t*_ and the multilocus inbreeding coefficient 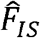 the estimate from both temporal samples (Weir & Cockerham, 1984). Selfing rate is assumed constant across generations. The method was implemented using DriftTest script (Navascués & Vitalis, 2020).

The *Osi* group was excluded from search for selection footprint given the small number of accessions available (70) and given the important disequilibrium in the number of accessions in Collect-1 and (15) and Collect-2 (55).

### Functional characterisation of loci under selection

Genes located within an interval of 250kb on both sides of independent loci detected to be under selection, were determined using the RAP-DB annotation of rice genome (https://rapdb.dna.affrc.go.jp/index.html). The corresponding MSU7.0 gene identity (TIGR) was retrieved using gene-ID converter tool proposed by RAP-DB. The window size of 250kb was chosen to represent approximately half the distance at which LD decayed to half its initial level, for both *Og* and *Osj* groups, estimated at 450kb on average across all chromosomes.

Enrichment analysis of genes within the chromosomic segments bearing loci under selection was performed using two gene ontology tools: Agri-GO (Tian et al., 2017; http://bioinfo.cau.edu.cn/agriGO/) and Panther-GO (Mi et al., 2019; http://GeneOntology.org), that use RAP and MSU gene identity, respectively, as input.

The chromosomic segments bearing loci under selection were also investigated for genes known to be involved in reproductive development processes, using available literature (Wei et al., 2020; http://www.modelcrop.org/). Likewise, chromosomic segment bearing loci under selection were investigated for enrichment in quantitative trait loci (QTL) involved in DTHD, using the Gramene QTL database (http://www.gramene.org/).

### Analysis of the temporal evolution of agro-climatic parameters

Climatological data (rainfall and temperature) of the 1961-2010 period were extracted from the archives of Guinea National Weather Service for a weather station in tropical forest area (N’Zérekoré, 7°48’53.2”N, 8°42’14.11”W) and a second station in the Sudanian savannah (Kankan, 10°23’01.65”N, 9°18’18.72”W) of the study region. Evolution of the annual rainfall and of the annual crops’ growth season rainfall was characterised using the Lamb index (Lamb, 1982): *I* = (*X*_*i*_ − *X*)/ *σ*, where *I* is the standardised anomaly (Lamb index), *X*_*i*_ is the climatic variable (rainfall) for the year *i* (or a portion of the year *i*), X is the average of the same variable in a reference period considered as normal, and *σ* is the standard deviation of the variable in the reference period. The normal meteorological reference period considered was 1961-1990, as recommended by the World Meteorological Organisation (WMO, 1996). The period considered as “crops’ growth season” was from June 1st to the end of October. The same methods were used to characterise the evolution trends of the minimum and maximum temperatures.

## Results

### Temporal evolution of major agro-climatic parameters in the study area

Patterns of deviation from the normal reference, of the annual rainfall total (ART) and of the annual rainfall total during the crops growing season (GSART) along the period 1961-2010, in our study area, are given in S Figure 2 and Figure 1, respectively. In both the Sudanian savannah area (Kankan) and the tropical forest area (N’Zérekoré), one can distinguish three major periods, though the beginning and the end of each of these periods are not exactly the same in the two areas. A rather humid period during the sixties, harboring positive deviations from normal, i.e., higher than normal rainfall. A dry period during the seventies and eighties, harboring mainly negative deviations from the normal. An instability period, from the beginning of the nineties until 2010, with contrasted year-to-year deviations from the normal rainfall. The seventies-eighties dry period is particularly marked in the Sudanian savannah area (Figure 1). A strong correlation was observed between ART and GSART, r = 0.953 in Kankan and r = 0.840 in N’Zérekoré.

**Figure 1:**
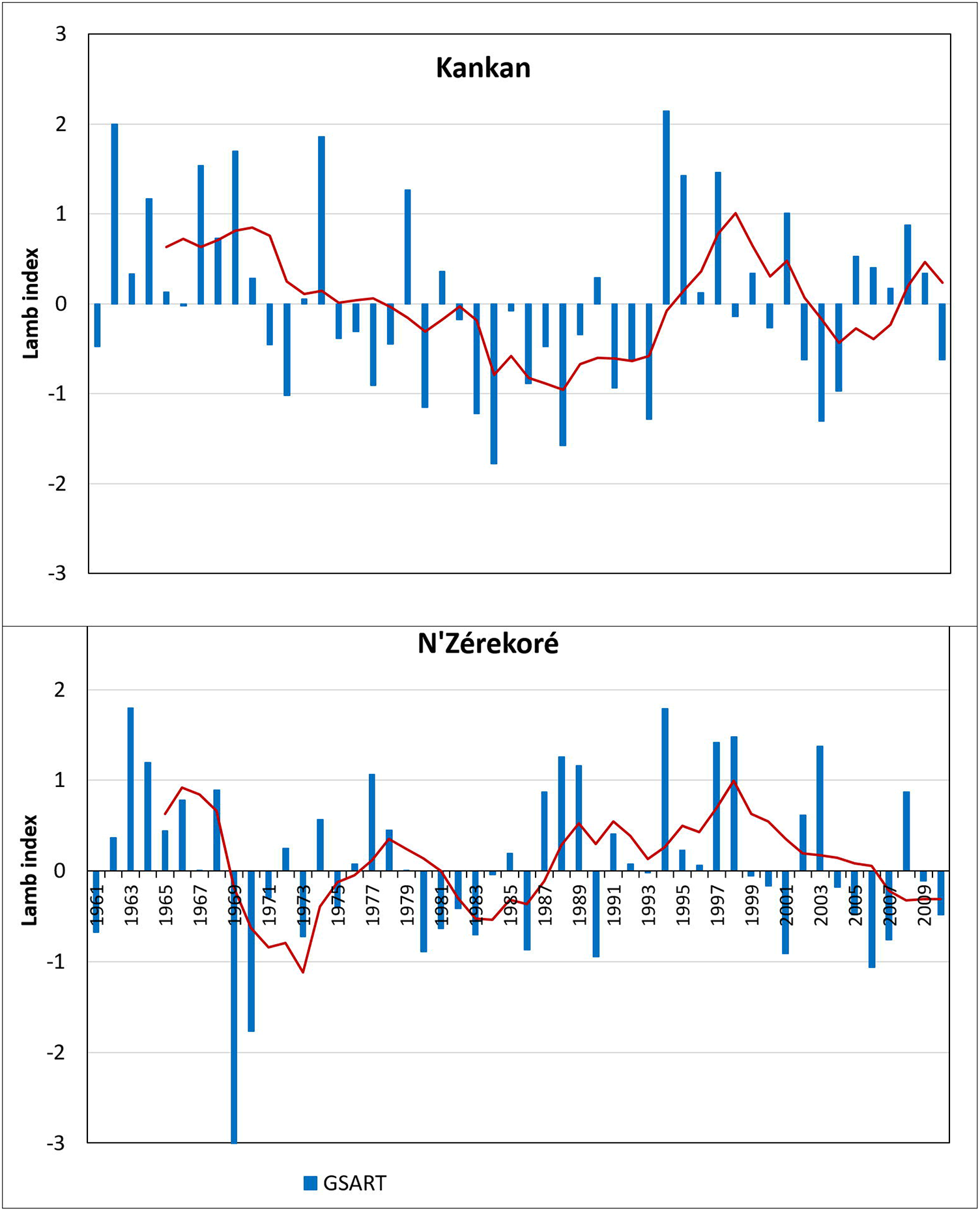
Pattern of deviation of the annual rainfall total during the crop growth season (GSART) from the normal reference, during the 1961-2010 period, in Kankan (10°23’01.65”N; 9°18’18.72” W) and N’Zérekoré (7°’48’53.2”N; 8°42’14,11”W) sites of Guinea.

Regarding the temperatures, patterns of the annual average minimum (T-Min) and maximum (T-Max) in the tropical forest area showed a systematic positive deviation from the normal reference from 1987 on (Figure 2). The average T-Min and T-Max of the period 1991-2010, were higher than the average T-Min and T-Max of the period 1961-1990, by 1.07°C and 0.65°C, respectively. In the Sudanian savannah area also, the annual average T-Max showed a systematic positive deviation from 1987 on, the average difference between the 1991-2010 and 1961-1990 periods being of 1.02°C. The annual average T-min showed less systematic deviation from the normal reference, and the average difference was -0.14°C (Figure 2).

**Figure 2:**
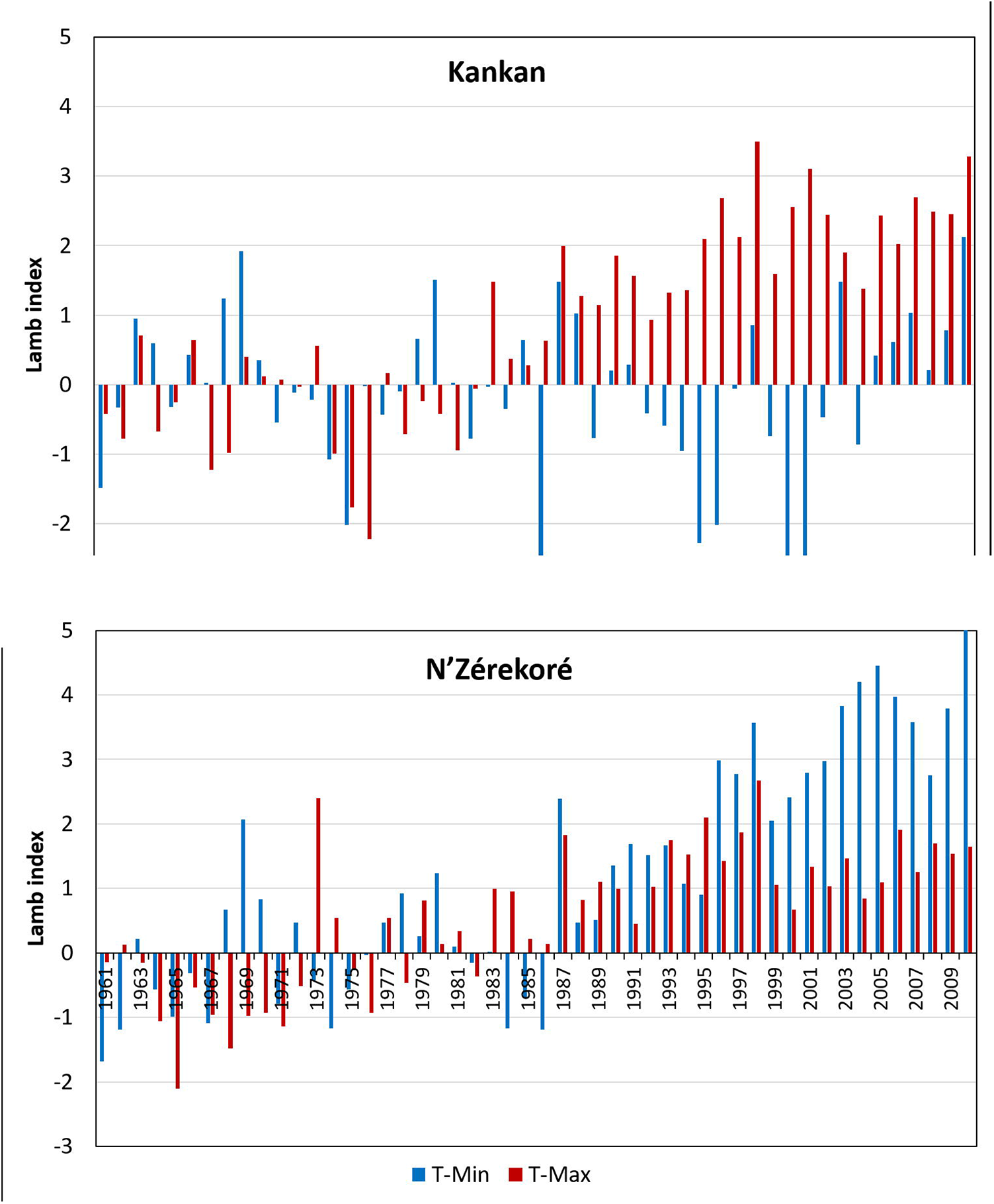
Pattern of deviation of the annual average minimum (T-Min) and maximum (T-Max) temperature from the normal reference, during the 1961-2010 period, in Kankan (10°23’01.65”N; 9°18’18.72” W) and N’Zérekoré (7°’48’53.2”N; 8°42’14,11”W) sites of Guinea.

### Variation in day to heading

A large variability of day to heading (DTHD) was observed among the 651 accessions phenotyped (77 < DTHD < 161 days) and between the three genetic groups (145, 115, and 106 days on average in *Og, Osi* and *Osj*, respectively). No significant relationship was found between the accessions’ DTHD and the latitudinal coordinate of their place of collect (R < 0.15) in both Collect-1 and Collect-2 subpopulations of each genetic group.

The distributions of the frequency of accessions of the two collect times in different classes of DTHD, are presented in Figure 3. These distributions show an important shift of frequencies of Collect-2 accessions toward shorter DTHD, for the genetic groups *Og* and *Osj*, and a more complex pattern for *Osi*. On average, the DTHD of *Og*-2 and *Osj*-2 were significantly shorter than the DTHD of their Collect-1 counterpart: 10.3 days between *Og*-1 and *Og*-2 (p < 0.00001) and 3.4 days between *Osj*-1 and *Osj*-2 (p < 0.002) (S Table 2). Conversely, the DTHD of *Osi*-2 was, on average, 4.3 days longer than the DTHD of *Osi*-1 accessions. The probability of significance of this average difference was rather low, p < 0.048 (S Table 2).

**Figure 3:**
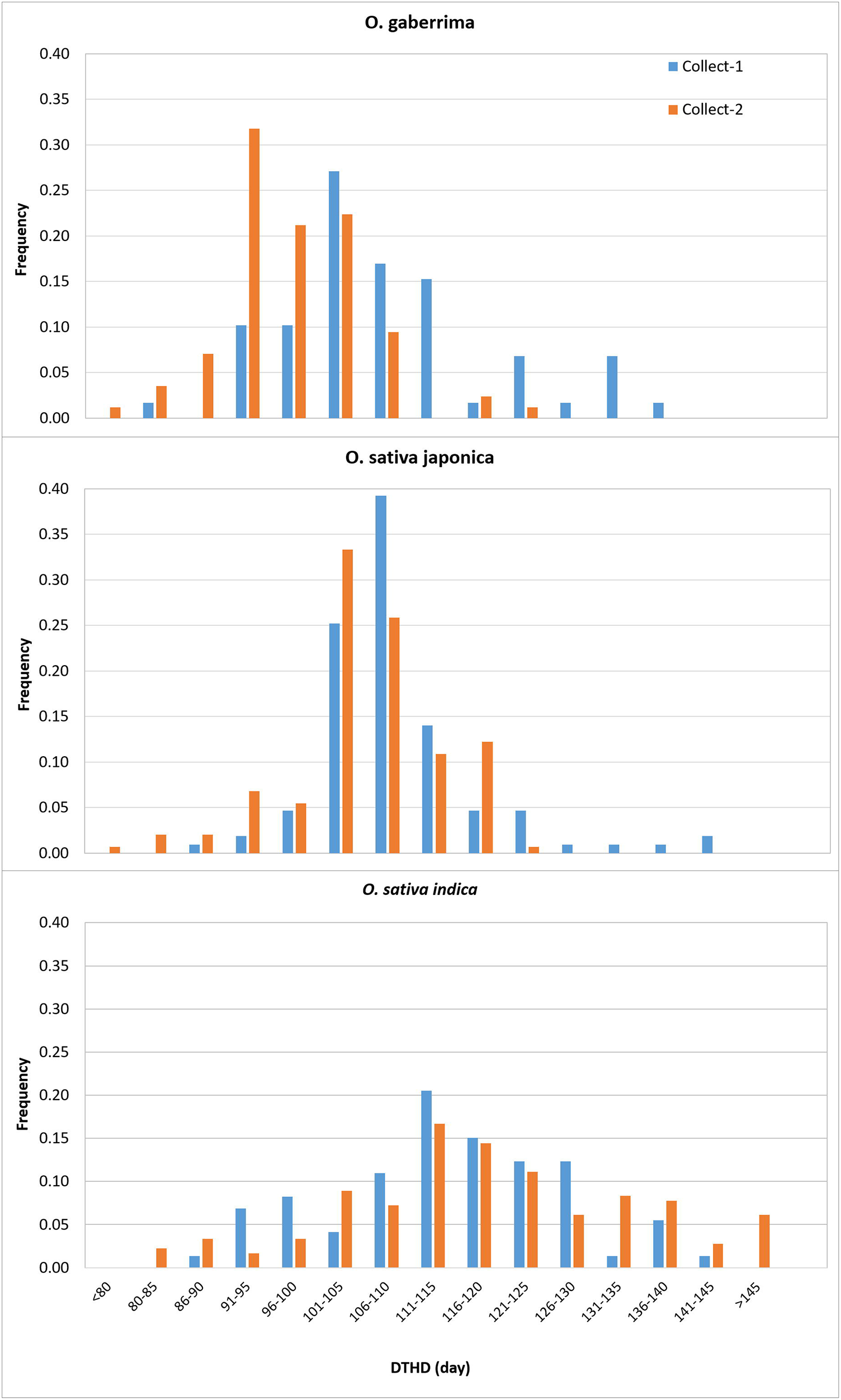
Distribution of frequency of accessions of the two collect times (Collect-1 and Collect-2) in different classes of duration of the vegetative phase expressed in number of days between the sowing date and the heading date (DTHD).

### Genetic diversity and population parameters

Pattern of genetic diversity drawn from genotypes at 1,130 SNP loci common to the 530 accessions genotyped with the GBS technology, showed three clear-cut clusters corresponding to the three well-known genetic groups: *Osi*, (*Osj*) and *Og* (Figure 4). Within each cluster, accessions from the two collect times did not show plausible preferential grouping (S Figure 3).

**Figure 4:**
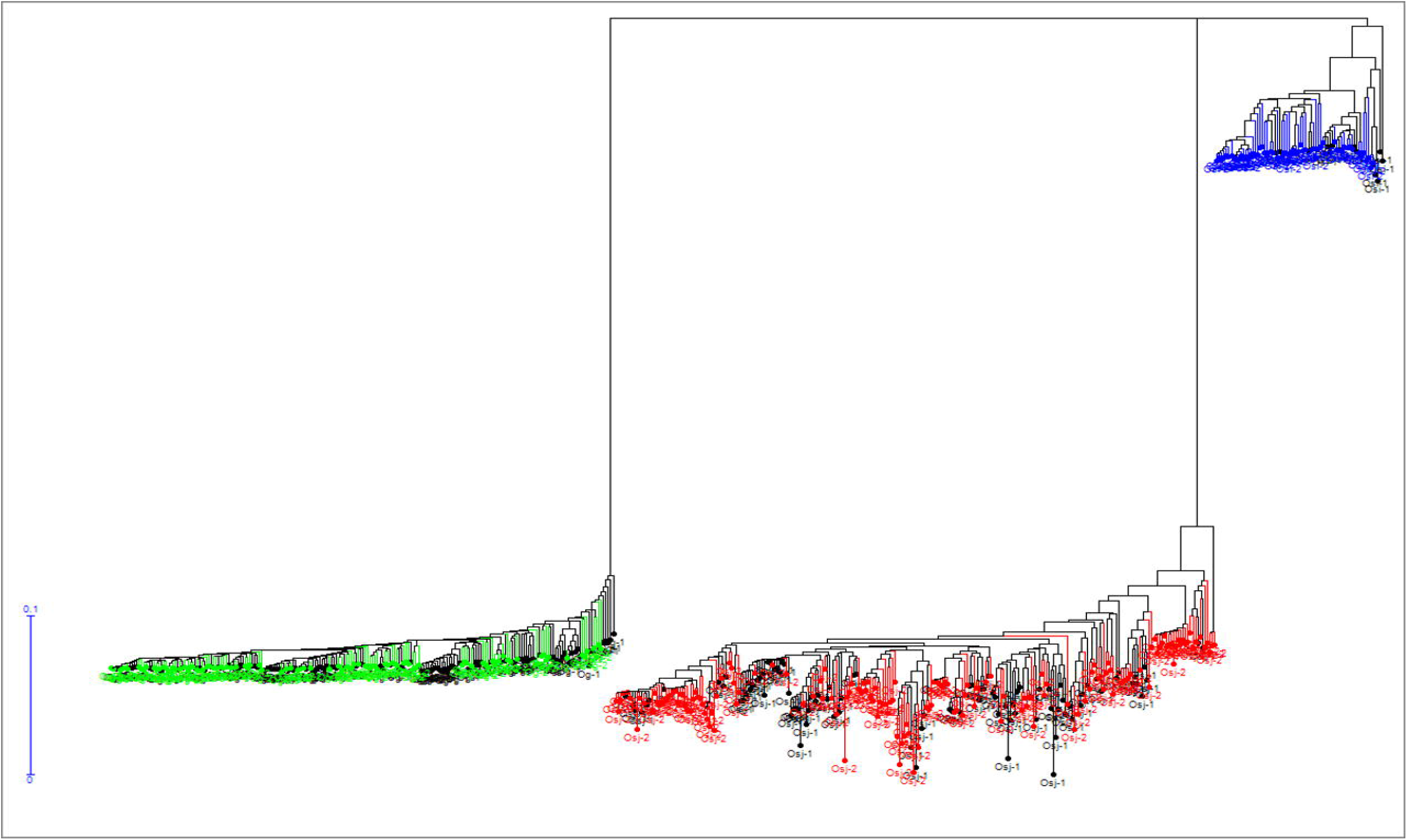
Unweighted neighbor-joining tree of simple matching distances constructed from genotypes of 530 rice accessions at 1,130 SNP loci. Tree branches bearing green, red and blue colors correspond to *O. glaberrima, O. sativa indica* and *O. sativa japonica* groups, respectively. Within each branch, accessions from the first collect time (Og-1, Osi-1 and Osj-1) are shown in black.

Nucleotide diversity was very low for *Og*-1 and *Og*-2 subpopulations (> 0.06), low for *Osj*-1 and *Osj*-2 (> 0.09), and of intermediate level (> 0.21) for *Osi*-1 and *Osi*-2 (S table 3). The genetic diversity estimated with the *θ* parameter showed a similar trend, with the lowest values for *Og*-1 and *Og*-2 (Mean *θ*_*π*_ = 319), the intermediate values for *Osj*-1 and *Osj*-2 (Mean *θ*_*π*_ = 798) and the highest values for *Osi*-1 and *Osi*-2 (Mean *θ*_*π*_ = 2,867) (S table 3). The Tajima’s D test detected significant deviations from neutrality in *Og*-1 and *Og*-2 (Mean D = -2.01, p-value = 0.0010), almost significant deviation in *Osj*-1 and *Osj*-2 (Mean D = - 1.29, p-value = 0.0575), and non-significant deviation in *Osi*-1 and *Osi*-2 (S table 3). The *F*_*IS*_ values ranged between 0.0001 (for *Osi*-1 and *Osi*-2) and 0.0303 for *Og*-2), suggesting a random union of gametes within each population.

Within each group the “among population” (i.e., among Collect-1 and Collect-2) molecular variance computed by AMOVA was extremely small (< 1%) and the “among individuals within populations” variance did not exceed 2.5%. The remaining large share of molecular variance was attributed to “within individuals” variance (S table 3). Genetic differentiation between the two subpopulations of each group was also very small (*F*_*ST*_ = 0.00014 between *Og*-1 and *Og*-2, *F*_*ST*_ = 0.00004 between *Osj*-1 and *Osj*-2, and *F*_*ST*_ < 0.00001 between *Osi*-1 and *Osi*-2), suggesting random distribution of genotypes across populations (S table 3).

Important differences were observed between groups for the r^2^ estimates of the initial LD, i.e. pairwise distance between markers below 25 kb: 0.7265 for *Osj*, 0.601 for *Og*, and 0.387 for *Osi*-2. The number of accessions in the *Osi*-1 subpopulation (15) was too small to provide a good estimate of LD. The three groups also differed for the subsequent decay of LD (Figure 5, S Table 4). While the distance at which the LD went below 0.2 was between 200 and 250 kb for *Osi*, it was between 500 and 700 kb for *Og*-1 and *Og*-2, and above 2,000 kb for *Osj*-1 and *Osj*-2. The speed of LD decay was slightly higher in *Og*-2 and *Osj*-2, compared to *Og*-1 and *Osj*-1, respectively.

**Figure 5:**
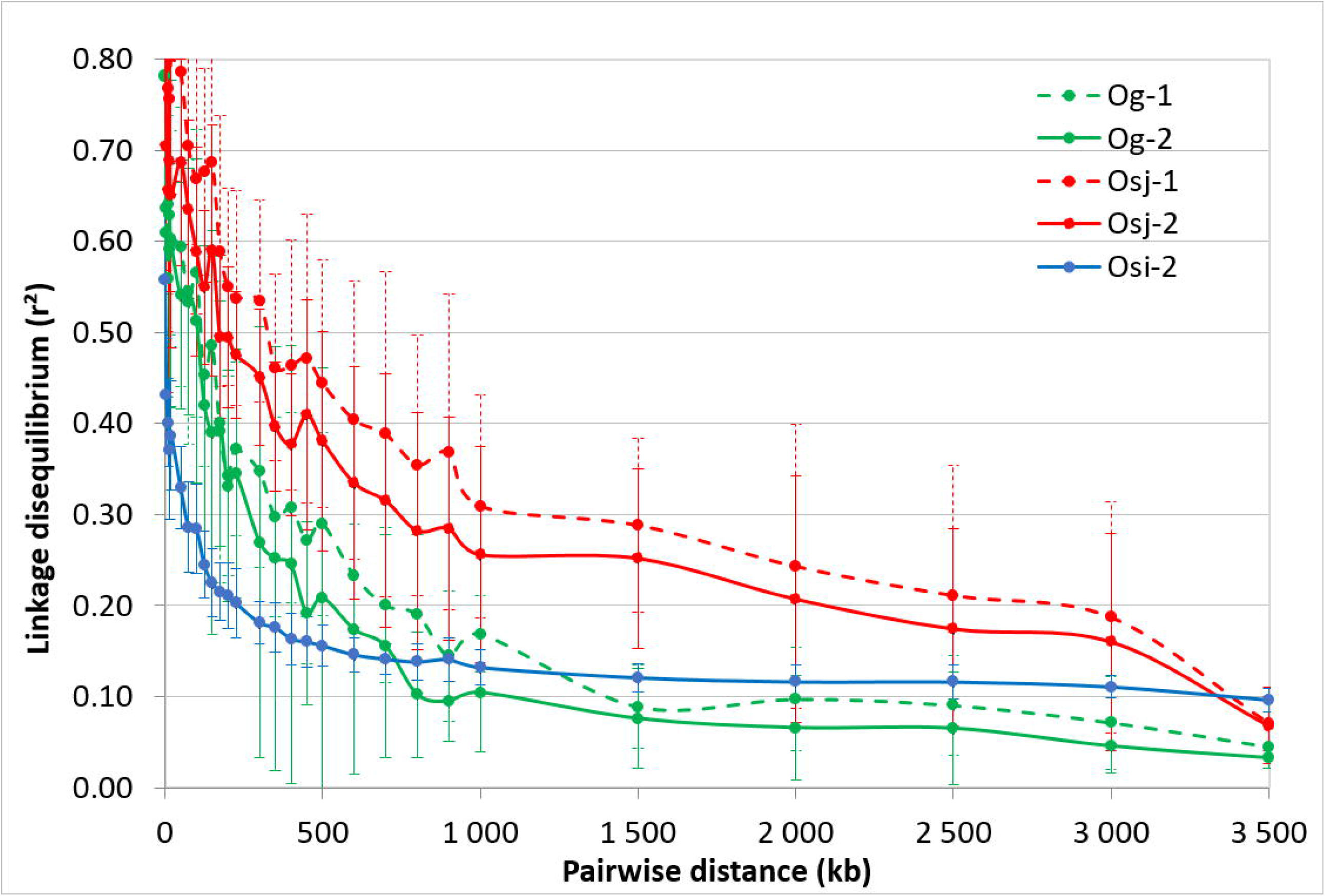
Patterns of decay in the r^2^ estimate of linkage disequilibrium (LD), in the rice populations collected in 1980 (xx-1) and in 2011 (xx-2); *Og*-1 and *Og*-2: *O. glaberrima*; *Osj*-1 and *Osj*-2: *O. sativa japonica*; Osi-2, *O. sativa indica*. The LD was estimated from 6,250 SNP, 9,282 SNP and 14,775 SNP loci for *Og, Osj* and *Osi* populations, respectively. The curve represents the average r^2^ according to pairwise distance between markers among the 12 chromosomes and the bars represent the associated standard deviation.

### Search for selection footprint

Search for selection footprint among the 6,250 SNP loci of the *Og* group, using the heterozygosity-based method, identified 20 SNP loci with *F*_*ST*_ value beyond the 1% confidence interval limits obtained from simulated data for *Og* group, and 54 additional SNP loci with *F*_*ST*_ value within the 1-5% confidence interval limits obtained from simulation (Figure 6). The former 20 SNP corresponded to 14 independent loci (*IL*), i.e. distance between adjacent SNP loci below 250 kb and strong LD (r^2^ > 0.2, often close to 1). Among the latter 54 SNP, 21 clustered with five of 14 previously defined *IL*. The remaining 33 SNP formed 16 additional independent loci composed of one to 14 SNP (S Table 5). Distances between each SNP under selection and its two adjacent SNP not under selection were 56.5 kb on average, did not exceed 363 kb, and, thus, were below the distance at which the LD exceeded the value r^2^ > 0.2. The LD between the SNP under selection belonging to different *IL* and different chromosomes were low (r^2^ < 0.1); it exceeded 0.2 in 13% cases out of the 242 possible combinations of *IL* but never reached 0.5 (S Figure 4). The heterozygosity in SNP loci under selection had fallen from 1.56% in the *Og-1* subpopulation to 0.76% in *Og-2* subpopulation.

**Figure 6:**
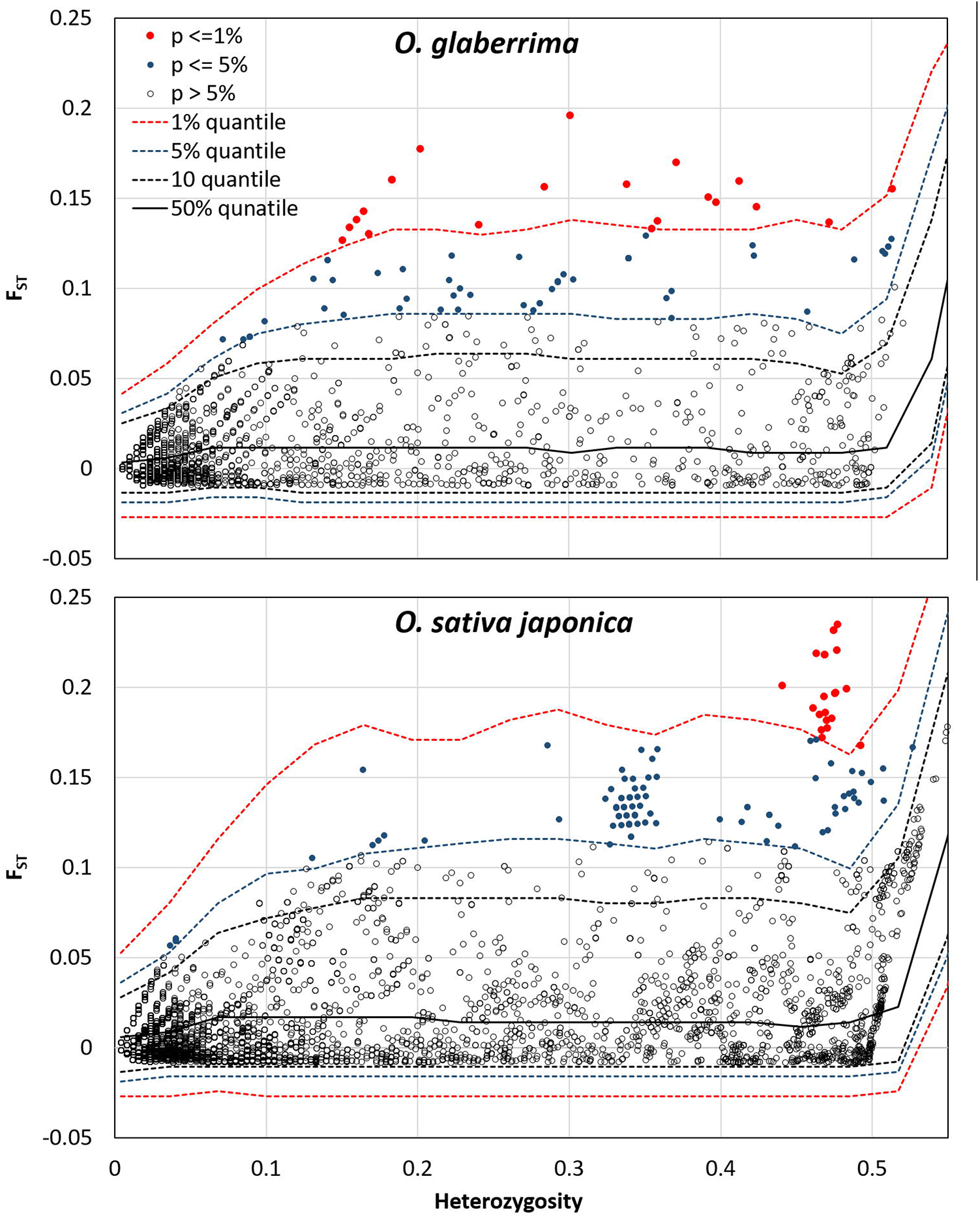
Plot of the joint distribution of F_ST_ and heterozygosity within populations for the observed loci (circles) and the one-sided confidence interval limits obtained from simulated data (dashed lines). Loci departing from neutrality hypothesis at 1% and 5% significance levels are indicated in red and blue circles, respectively. The number of observed loci was 6,250 for *O. glaberrima*, 9,282 for *O. sativa japonica*.

Search for selection footprint among the 9,282 SNP loci of *Osj* group yielded 22 SNP loci with *F*_ST_ value beyond the 1% confidence interval limits obtained from simulated data for *Osj*, and 88 additional SNP loci with *F*_S*T*_ value within the 1-5% confidence interval limits obtained from simulation (Figure 6). The former 22 SNP corresponded to three *IL*. Among the latter 88 SNP, 16 clustered with the three previously defined *IL*. The remaining 72 SNP formed 11 additional *IL*, including a cluster of 44 SNP (S Table 5). Distance between each SNP under selection and its two adjacent SNP not under selection was 49.6 kb on average, did not exceed 270 kb and, thus, were well below the distance at which the LD exceeded the value r^2^ > 0.2. The LD between the SNP under selection belonging to different *IL* and different chromosomes was low (r^2^ < 0.1); the r^2^ value exceeded 0.2 in 17% cases out of the 125 possible combinations of independent loci and exceeded 0.5 in two combinations (S Figure 4). The heterozygosity in SNP loci under selection had fallen from 1.77% in the *Osj-1* subpopulation to 1.02% in *Osj-2* subpopulation.

Search for selection footprint using the drift-based approach, among the 6,250 *Og* SNP loci, identified 29 SNP with *F*_*ST*_ departing from the neutrality hypothesis with q-value < 0.05 (Table 2; S Table 5). All these loci were also detected to be significantly under selection with P <= 0.01 (19 loci) or P<= 0.05 (10 loci) with the heterozygosity-based detection method. Likewise, among the *Osj* SNP loci, 30 showed *F*_*ST*_ with q-value < 0.05 (S Table 5). All these loci were also detected to be significantly under selection with P <= 0.01 (13 loci) or P<= 0.05 (17 loci) with the heterozygosity-based detection method. Thus, the results of the drift-based method corroborate the results of heterozygosity-based method, especially for loci most significantly under selection.

**Table 2:**
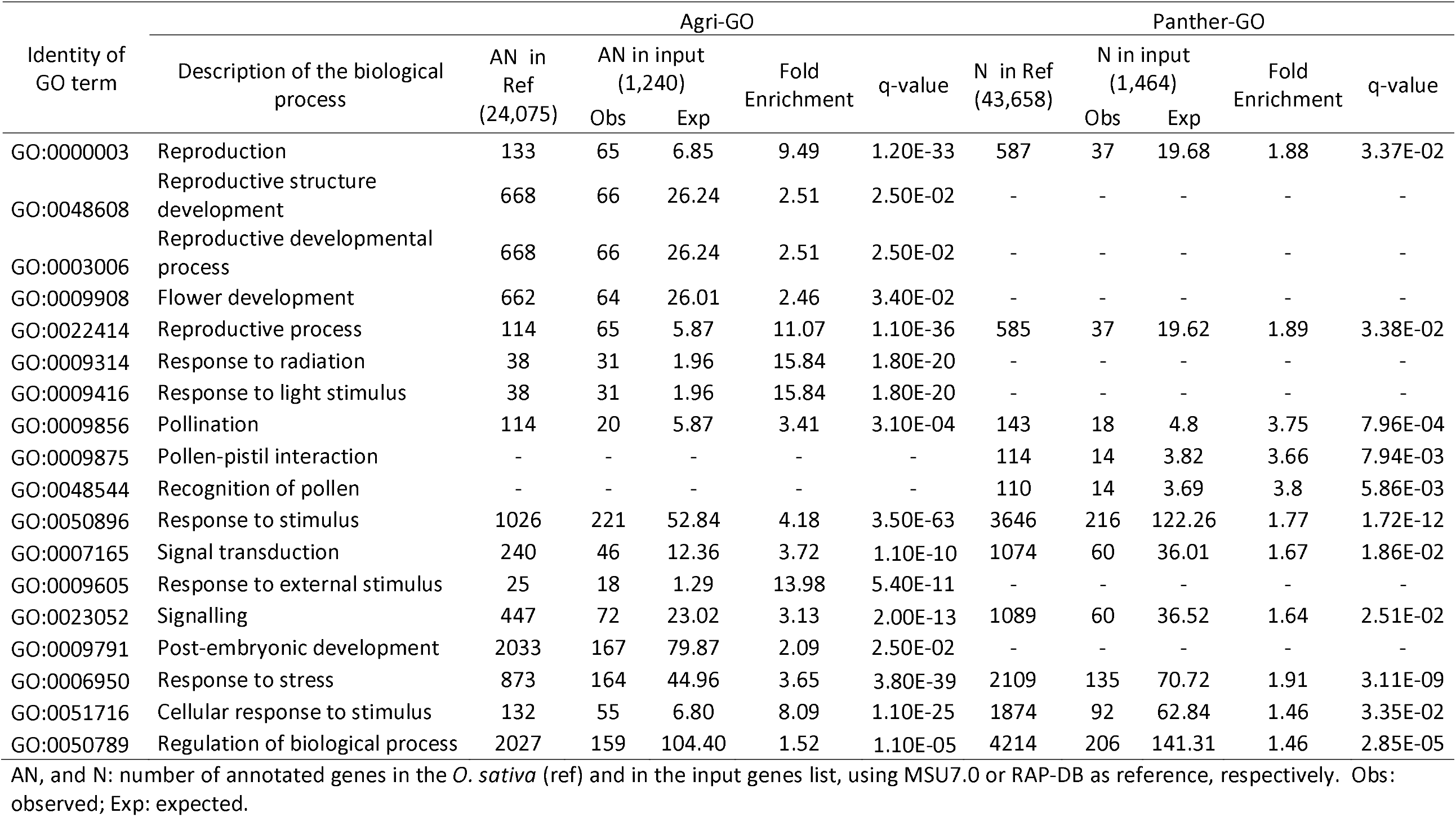
Enrichment in gene ontology terms related to reproductive biological processes, of the chromosomic segments bearing the 24 independent loci most likely under section.

### Functional characterisation of loci under selection

Prior to functional characterisation, six independent loci under selection (*ILUS*) in *Og* and in *Osj*, belonging to the same chromosomes and within distance between the component SNP below 250kb, were merged. The process reduced the total number of *ILUS* to 40, each composed of one to 45 individual SNP. Among these *ILUS*, nine, composed of one SNP and identified to be significantly under selection by one detection method only, were discarded from further analyses. The remaining 31 *ILUS* were divided into two categories: *ILUS* including at least one SNP identified to be significantly under selection by the two detection methods (24 cases, assembling 139 SNP, hereafter referred to as “M2 type”) and *ILUS* detected by one method only, namely the heterozygosity-based method (7 cases, assembling 36 SNP), hereafter referred to as “M1 type”.

For functional characterisation purpose, the chromosomic interval of each *ILUS* was extended by 250 kb at its two extremities to account for the extent of LD. When concatenated, the 24 M2 type *ILUS* represented a pseudo-chromosomic segment of 23.26 Mb length, bearing 2 038 genes endowed with an MSU-TIGR gene identity or 1,517 genes endowed with RAP gene identity. The addition of the seven M1 type *ILUS* stretched the length of the pseudo-chromosomic segment to 30.77 Mb, bearing 2,706. MSU-TIGER genes or 2,612 RAP-DB genes.

The ontology analysis of the 2,038 MSU-TIGR genes, on the pseudo-chromosomic segment bearing the 24 M2 type *ILUS*, with the Agri-GO tool, detected highly significant enrichment in genes involved in several biological processes (S Table 6A) including “reproductive processes”, “response to light and radiation” and “response to stimulus” (Table 2). Shift to 2,706 MSU-TIGER genes (i.e. inclusion of 668 genes underlining the seven M1 type *ILUS*) did not markedly modify the pattern of gene enrichment (S Table 6B). The ontology analysis of the 1,517 RAP-DB genes, using Panther-GO tool revealed an enrichment pattern (S Table 6C) that included an important share of the ontology terms related to reproductive processes previously detected with Agri-GO, plus two ontology terms related to biological processes of “recognition of pollen” and “pollen-pistil interaction” (Table 2). Shift to 2,612 RAP-DB genes did not markedly modify the pattern of gene enrichment (S Table 6D).

Investigation of the list of genes underlying the 24 M2 type *ILUS* for genes reported to be involved in reproductive processes, identified 12 genes (Table 3, S Table 7) including OsGI (“Gigantea”) related to the circadian clock, HD1 (“Constans”) involved in induction of flowering under inductive short days, and OsPHYB (Phytochrome B) involved in the regulation of critical day length for flowering. When all the 31 *ILUS* were considered, three additional known genes involved in biological processes related to heading were identified (Table 3, S Table 7).

**Table 3:** Genes and QTLs located on the chromosomic segments bearing the independent loci under selection.

| Independent loci under selection |  |  |  |  |  | Genetic group involved | DTHD QTLs (2) |  |  | DTHD genes (3) |  |  |
| --- | --- | --- | --- | --- | --- | --- | --- | --- | --- | --- | --- | --- |
| N° | Type | Nb of SNP (1) | Chr | Start | End |  | Nb | Known name | MSU Identity | Name | GRP | Reference |
| 1 | M2 | 2 | chr01 | 3 930 523 | 4 504 261 | *Og & Osj* | 1 |  | LOC_Os01g08700 | OsGI | 1 | Izawa et al. (2011) |
| 2 | M2 | 1 | chr01 | 5 540 432 | 6 040 432 | *Og* | 14 | dth1.1, QHd1a, QHd1a | LOC_Os01g10590 | OsFTL8 | 1 | Zhang et al. (2020) |
| 3 | M2 | 9 | chr01 | 18 423 679 | 19 143 461 | *Og* | 2 |  |  |  |  |  |
| 4 | M1 | 2 | chr02 | 6 762 905 | 7 267 332 | *Osj* | 4 |  |  |  |  |  |
| 5 | M2 | 1 | chr02 | 24 604 636 | 25 104 636 | *Og* | 12 | qHD-2, QHd2a | LOC_Os02g41550 | OsCRY2 | 1 | Hirose et al. (2006) |
| 6 | M2 | 1 | chr03 | 5 079 703 | 5 579 703 | *Og* | 15 | dth3.1, QHd3b |  |  |  |  |
| 7 | M2 | 3 | chr03 | 10 132 774 | 12 145 360 | *Og* | 6 | Hd8, dth3.1 | LOC_Os03g19480 | OsSWN; SDG718 | 2 | Liu et al. (2014) |
| 7 | M2 | 3 | chr03 | 10 132 774 | 12 145 360 | *Og* | 6 | Hd8, dth3.1 | LOC_Os03g19590 | OsPHYB | 2 | Ishikawa et al. (2011) |
| 8 | M1 | 4 | chr04 | 0 | 827 020 | *Og & Osj* | 1 | qDTH-4 |  |  |  |  |
| 9 | M1 | 9 | chr04 | 1 085 381 | 3 005 652 | *Osj* | 12 | qDTH-4, QHd4 |  |  |  |  |
| 10 | M2 | 4 | chr04 | 3 993 589 | 4 670 203 | *Og & Osj* | 0 |  | LOC_Os04g08034 | OsEMF2 | 3 | Luo et al. (2009) |
| 11 | M2 | 1 | chr04 | 32 129 645 | 32 629 645 | *Og* | 2 | dth4.2 |  |  |  |  |
| 12 | M2 | 6 | chr05 | 8 974 371 | 9 788 465 | *Og* | 2 |  |  |  |  |  |
| 13 | M2 | 15 | chr05 | 14 743 007 | 16 632 418 | *Osj* | 2 | qHD-5 |  |  |  |  |
| 14 | M2 | 8 | chr06 | 9 422 923 | 10 942 740 | *Osj* | 3 | Hd1 | LOC_Os06g16370 | Hd1 | 1 |  |
| 14 | M2 | 8 | chr06 | 9 422 923 | 10 942 740 | *Osj* | 3 |  | LOC_Os06g16390 | OsCLF; SDG711 | 2 | Liu et al. (2014) |
| 15 | M2 | 1 | chr06 | 11 153 613 | 11 653 613 | *Osj* | 0 |  |  |  |  |  |
| 16 | M2 | 45 | chr06 | 13 095 417 | 18 074 177 | *Og & Osj* | 5 | Hd6b, dth6.1 | LOC_Os06g30370 | OsMFT1 | 1 | Gao et al. (2014) |
| 17 | M2 | 1 | chr07 | 17 212 559 | 17 712 559 | *Og* | 5 |  |  |  |  |  |
| 18 | M2 | 1 | chr07 | 29 373 622 | 29 873 622 | *Og* | 6 | Hd2, QTL7c | LOC_Os07g49460 | OsPRR37 | 1 | Hu et al. (2021) |
| 19 | M2 | 2 | chr08 | 4 860 945 | 5 579 786 | *Osj* | 3 | Hd-5, dth8 |  |  |  |  |
| 20 | M2 | 14 | chr08 | 17 288 523 | 18 502 122 | *Osj* | 21 | QHd8, aQHd8, qDTH-8 |  |  |  |  |
| 21 | M1 | 2 | chr08 | 19 800 834 | 20 305 131 | *Og* | 2 | qDTH-8 |  |  |  |  |
| 22 | M2 | 1 | chr08 | 23 308 242 | 23 808 242 | *Og* | 2 | qDTH-8 | LOC_Os08g37490 | OsGF14a | 3 | Liu et al. (2019) |
| 23 | M2 | 1 | chr09 | 14 585 036 | 15 085 036 | *Og* | 3 | QHd9, FLTQ3 |  |  |  |  |
| 24 | M1 | 2 | chr09 | 19 597 853 | 21 078 706 | *Og* & *Osj* | 18 | qHDD9-1, qDTH-9 | LOC_Os09g33850 | OsFTL4 | 3 | Fang et al. (2019) |
| 24 | M1 | 2 | chr09 | 19 597 853 | 21 078 706 | *Og* & *Osj* | 18 | qHDD9-1, qDTH-9 | LOC_Os09g34070 | OsFPA | 2 | Mouradov et al. (2002) |
| 24 | M1 | 2 | chr09 | 19 597 853 | 21 078 706 | *Og* & *Osj* | 18 | qHDD9-1, qDTH-9 | LOC_Os09g36220 | OsPRR95 | 1 | Murakami et al. (2005) |
| 25 | M2 | 2 | chr09 | 22 453 022 | 22 953 023 | *Og* | 1 | qDTH-9 |  |  |  |  |
| 26 | M1 | 3 | chr11 | 3 400 294 | 3 922 487 | *Osj* | 1 | dth11.1 |  |  |  |  |
| 27 | M2 | 1 | chr11 | 23 361 713 | 23 861 713 | *Osj* | 2 | QHd11 |  |  |  |  |
| 28 | M2 | 14 | chr11 | 25 384 918 | 26 543 152 | *Og* & *Osj* | 1 | hd11 |  |  |  |  |
| 29 | M2 | 4 | chr11 | 26 942 125 | 27 922 719 | *Og* | 0 |  |  |  |  |  |
| 30 | M1 | 14 | chr12 | 16 142 460 | 17 897 080 | *Og* | 12 | dth12.1, QHd12 |  |  |  |  |
| 31 | M2 | 1 | chr12 | 20 923 730 | 21 423 730 | *Og* | 4 |  | LOC_Os12g34850.1 | OsVRN5; OsVIL2 | 2 | Wang et al. (2013) |

RP: heading time regulation pathway. (1) Photoperiod and circadian clock pathway, (2) chromatin-related pathway, and (3) hormones pathway. Type: M1= Independent loci under selection detected under one detection method; M2= Independent loci under selection including at least one locus detected to be under selection by the two detection methods implemented. *Og*: *O. glaberrima*; *Osj*: *O. sativa japonica* (1) Number of SNP loci under selection composing the independent loci; chr: Chromosome; DTHD: day to heading; (2) source http://www.gramene.org/; (3) see detailed data and references in S Table 7.

Investigation of the chromosomic segments bearing the 24 M2 type *ILUS* for the presence of QTLs involved in DTHD, counted 112 such QTL out of the 619 listed in the Gramene QTL database (http://www.gramene.org/) (Table 3; S Table 7). This represented a density of one DTHD QTL every 0.21 Mb, approximately 3.0-fold higher than the expected average density of DTHD QTL, one every 0.62 Mb. The 112 DTHD-QTL counted included a number of major QTL such as Hd1, Hd2, Hd5, Hd6, Hd8 and Hd11. When all the 31 *ILUS* were considered, the count of DTHD QTL reached 162, representing a 3.3-fold enrichment.

## Discussion

Understanding changes in the phenotypic and genetic diversity of crop species undergoing environmental selection pressure can provide a wealth of information regarding the genetic processes at work and the adaptation strategies (Snowdon et al., 2021). This study provided insight into the extent of the climate-changes, the related selection pressure on meta-populations of a mainly autogamous crop species, rice, into the extent of its adaptive phenological response, and into the underlying adaptive genetic processes, in the context of a rapid adaptation over 30 generations.

The extent of changes in climate parameters, temperatures and rainfall, between our two collect times (1980 and 2011), were similar to that reported for the entire West Africa (Sultan et al., 2019). It was characterized by approximately a 1 °C rise in mean T-Max and mean T-Min, and by a severely dry period followed by a period of high variability of the annual and cropping season rainfalls. Analysing N’Zérekoré rainfall data for the 1931-2014, Loua et al. (2017) reported that the higher variability of the annual rainfall during the period 1994 – 2014 was associated with a shorter duration of the rainy season (later beginning and earlier end) and a less predictable date for the establishment of the rainy season.

The extent of the heading date advance (10.3 days on average) in our *Og-2* subpopulation, compared to *Og-1*, was among the highest reported for other plant species in relation with climate changes. For instance, the flowering time advancement was of seven days on average, for 43 common plant species of North America for a temperature rise of 2.5 °C, over some 150 years (Miller-Rushing, 2008). It was of two days in a 22-generation resurrection experiment of the model legume *Medicago truncatula* (Gay et al., 2021), and of 5 to 10 days °C^-1^, according to ecotypes, in *Arabidopsis thaliana*, under experimental conditions (Footitt et al., 2021). In comparison, the advance in DTHD of the *O. sativa* subpopulation *Osj-2* relative to *Osj-1* (3 days on average) was on the lowest side of the reported flowering time advancement. Important plasticity of the heading date has been reported in rice, in relation with the temperatures during the vegetative growth (Guo et al., 2020). However, the “temporal clean” setting of our study, (phenotyping of samples of the two collect times under the same environmental conditions) excludes the role of this adaptive mechanism in the advanced DTHD of *Og-2* and *Osj-2* subpopulations compared to *Og-1* and *Osj-1*, respectively. On the other hand, an increase in temperatures, within the range of the observed 1°C, mainly affects rice grain yield due to an increase in maintenance respiration and to a decrease in grain formation (reviewed in Wassmann et al., 2010). Thus, the very significant advance of DHTD in *Og-2* and *Osj-2*, compared to their *Og-1* and *Osj-1* counterpart, most probably results from changes in rainfall regime. This hypothesis is also supported by the fact that the shorter DTHD were observed in *Og-2* and *Osj-2* subpopulations almost exclusively cultivated in the drought-prone upland or hydromorphic ecosystems, and not in the *Osi-2* subpopulation, cultivated in the lowland ecosystem, less prone to the effects of changes in rainfall regime. The very low coefficient of determination between the latitudinal coordinate of the collect place and the length of DTHD, whatever the collect time and the subpopulation, suggests that the environmental selective pressure was homogenous across our study areas stretching between 7°34’19.96”N and 12°29’1.50”N.

Literature describing plant response to climate changes often refers to natural populations and distinguishes three types of responses: migration, temporary (reversible) adaptation through phenotypic plasticity and less reversible genetic adaptation (Franks & Hoffmann, 2012; Gray & Brady, 2016). When dealing with crop-species populations, one should also consider human (farmers) actions (selection) that can accelerate or slow down plant populations’ natural responses. Farmers’ actions can take the drastic form of replacement of the existing crop varieties or the more subtle form of selection among the standing diversity or among new diversity emerging from the mutation and/or recombination processes.

In our case, several facts argue against the hypothesis of replacement of the old rice landraces of long DTHD (*Og-1* and *Osj-1* subpopulations) by new exogenous rice varieties of shorter DTHD (*Og-2* and *Osj-2* subpopulations). (i) Penetration of improved *O. sativa* rice varieties, with shorter duration, in the slash-and-burn based upland rice cropping system of our study area was reported to be extremely limited (Barry et al., 2009) and no breeding program exists for the improvement and release of new *O. glaberrima* varieties. (ii) Distance based neighbour-joining tree did not show any structuring in relation with collect time. (iii) Genetic differentiation between *Og-1* and *Og-2* and between *Osj-1* and *Osj-2* was very low (*F*_*ST*_ < 0.001). (iv) The genetic diversity (nucleotide diversity and *θ* parameter) of the Collect-1 and Collect-2 subpopulations had the same order of magnitude. The two latter observations also argue for the fact that the combined human and environmental selection pressure has not given rise to the emergence of a few well-adapted individuals (i.e., shorter DTHD) colonizing the entire area of cultivation of *O. glaberrima* and *O. sativa japonica*. A more subtle selection/genetic adaptation process seems to have been at work, preserving the overall neutral genetic diversity.

According to the theory of genetic adaptation (Orr, 2005), when a population is facing a novel environment, its potential for adaptive evolution is provided by the already present “standing genetic variation” or by *de novo* genetic variation arising during that bout of adaptation. In our case, while the occurrence of adaptive mutations cannot be excluded, their spread to a significant share of the *O. sativa* and *O. glaberrima* subpopulations is unlikely given the length of generation in rice (one year), the rather short interval (31 years) separating our two collect times and the rather low rate of outcrossing in rice. Indeed, the highest natural outcrossing rate reported is 6.8% in *O. sativa* (Sahadevan & Namboodiri, 1963) and 5% in *O. glaberrima* (Oka & Morishima, 1967). Analysing the distribution of rice genetic diversity at farm, village and ecosystem levels in Guinea, using SSR molecular markers, Barry et al. (2007b; 2007c) reported that (i) rice landraces had a multi-line genetic structure; (ii) the within- and between-farm *F*_*ST*_ values were of the same order of magnitude; (iii) within-farm genetic diversity was high, i.e., up to 50% of total genetic diversity observed at the village level; (iv) each village pooled more than half of the regional allelic diversity; (v) regional allelic diversity was comparable to that noted worldwide for the Asian rice (O. sativa), but not as high for the African rice (*O. glaberrima*). Thus, one can consider that the *O. sativa* and *O. glaberrima* rice populations of our study area had had the adaptive evolution potential needed to face the environmental and human selection pressure over the last 30 years. More generally, one can assume that *Og-1* – *Og-2* and *Osj-1* – *Osj-2* subpopulations are independent replicates of the evolutionary process, where migration had a negligible role.

Several methods have been proposed to investigate the role of selection, versus drift, in phenotypic changes between populations (Hansen et al., 2012). The temporal clean approach (Lande 1977; Goldringer & Bataillon, 2004) we have implemented offers the advantage of using plant material in which the populations have actually adapted in contemporary times to changes in climate. However, the majority of the statistical methods detecting selection footprint were developed for outcrossing populations, and assumes random mating, making challenging the detection of loci under selection in predominantly selfing populations (Hartfield et al., 2017; Navascués et al., 2021). Indeed, given the extent of LD, selective sweeps in selfing populations can involve large genomic segments and hamper distinction of the genetic features between neutral and adaptive loci. Moreover, selfing can limit adaptation by slowing down pollen-based flow of beneficial variations, can increase drift by reducing the number of independent alleles sampled at reproduction, and can affect the dynamic of polygenic adaptation (reviewed in Hartfirld et al., 2017). On the other hand, selfing can facilitate the detection of selective sweeps by reducing (i) the fixation times of beneficial alleles, exposing them faster to selection pressure (Charlesworth, 1992) and (ii) the number of new effective recombinations, making it easier to detect sweeps (Smith & Haigh, 1974). We implemented two genome scan methods, one assuming random mating (Excoffier et al., 2009) and another that introduces several modifications to the method proposed by Goldringer & Bataillon (2004) to take into account partial selfing (Navascués et al., 2021). The two methods identified several independent loci under selection, in both *O. glaberrima* (25 independent loci distributed on 11 chromosomes) and *O. sativa* (18, distributed on 8 chromosomes) groups, and all the loci, identified by the latter method to be under selection, were also identified as such by the former method, assuming random mating. These results suggest that gene flow and recombination were not the main genetic processes underlying the observed phenotypic adaptation, i.e. earlier flowering time. In other words, the outlier values of change in allele frequency in some loci, detected from the comparison of the observed and expected temporal genetic differentiation (*F*_*ST*_), were mainly the result of selection among the pre-existing allelic combinations. The above-described multiline genetic structure of rice landraces and the rather large and imbricated genetic diversity of rice at the farm, village and region levels (Barry et al., 2007b; 2007c) supports this hypothesis. In a given village, the multiline composition of a given landrace varies between farms, and the landrace can be considered as a meta-population the components of which are subject to farm-to-farm customary exchanges (Barry et al., 2007c). The same imbricated picture can be applied to groups of adjacent villages and farther. This setting of genetic diversity allows the environmental selection pressure to operate either directly (individuals plant/line with long DTHD fails to complete their life cycle when the cropping season ends earlier) or indirectly through farmers’ selection of panicles bearing well-filled grains that will serve as seed for the next generation. Indeed, hand-picking of individual panicles is the most widely shared harvesting practice, in the slash-and-burn itinerant rice cropping system of our study area.

The 31 independent loci detected to be under selection were scattered among 11 chromosomes out of the 12 that compose the rice genome. Their length did not exceed 1,500 kb (560 kb on average), save one case on chromosome 6 of *Osj*, composed of 45 significant SNP and extending over 4,478 kb. Distance between these loci and their two adjacent SNP not under selection was around 50 kb on average and seldom went beyond the values at which the LD exceeded r^2^ = 0.2. This almost absence of selective sweeps confirms the presence of the adaptive variants on multiple genetic backgrounds, well before the time *Og* and *Osj* populations undergone environmental selective pressure. The selective pressure has halved the heterozygosity rate of the loci under selection, already highly homozygous (> 98%), like all other loci not under selection. This adaptive process resemble the “evolutionary rescue” process in selfing plants described by Uecker (2017). However, in the case of *Og* and *Osj* populations, the variants were already, largely, in homozygous state, and the number of independent loci involved was quite high, 25 in *Og* and 18 in *Osj*.

Among the *ILUS*, a relatively small share (13% in *Og* and 17% in *Osj*) were in LD above r^2^ > 0.2. This limited long-distance LD between the independent loci, on each chromosome and across chromosomes, testifies that though polygenic, the adaptive process involved in *Og* and *Osj* did not translate into predominance of multilocus genotypes, as it is often the case in selfing plants (Hartfield et al., 2017; Gay et al., 2021). Though rare in selfing populations, such a pattern of genetic basis of adaptation is in accordance with the complex polygenic genetic basis of the phenotypic adaptation involved, the DTHD. Indeed, the 31 *ILUS* showed significant enrichment in genes involved in the reproduction processes, and 29 of the loci could be connected to QTLs or genes (or both) involved in DHTD.

Rice is a short-day plant in which the heading date is regulated by a network of genetic components that can be organized into at least three pathways: photoperiod and circadian clock, chromatin-related pathway, and hormonal pathway (reviewed in Wei et al., 2020). Representatives of each of these pathways are present among the 15 DHTD genes we identified to be located on chromosome segments under selection (Table 3). For instance, *OsGI*, the orthologue of *GIGANTEA* in *A. thaliana*, is strongly involved in the circadian clock system and *Hd1*, the orthologue of *CONSTANCE* in *A. thaliana*, is a major photosensitivity gene in rice. Over-expression of *GI* triggers higher expression of *Hd1* and rapid flowering under any day length (Wei et al., 2020). *SDG711* and *SDG718* genes, involved in the chromatin pathway, also regulate the expression of *Hd1* and, thus, flowering in short days (Liu et al., 2014). *OsphyB* regulates the *Hd1*-mediated expression of the rice Florigen *Hd3a* and critical day length; its function in floral induction is not affected by the photoperiod (Takano et al., 2005; Ishikawa et al., 2011). *OsEMF2* (embryonic flower) gene belongs to the polycomb group proteins that play important roles in the epigenetic regulation of gene expression. Alteration of *OsEMF2* has pleiotropic phenotypic consequences including early flowering time (and even skipping the vegetative phase) even under long-day conditions, and abnormal flower organs (Luo et al., 2009). *OsGF14* is strongly induced by soil-drought stress and triggers the abscisic acid-dependent response pathway (Liu et al., 2019). It also acts as a negative regulator of flowering by interacting with *Hd3a* (Purwestri et al., 2009). Thus, the most probable underlying genetic bases of the observed DTHD responses to climate changes are subtle changes in the genetic network regulating rice DTHD and not the drastic turning off/on of one major gene. Given the number of genes involved, it would be difficult to extract the effects of individual genes, especially as the changes in the genetic network most probably vary from one rice landrace to another according to farms and villages.

The genetic bases of plant adaptations to environmental changes are often described in terms of soft and hard sweeps. In a mutation-limited world, recent adaptation leads to hard sweeps and leaves clear and well-understood footprints in genomic diversity (Hermisson & Pennings, 2017). The complex footprints of selection due to climate changes (higher temperature, reduced rainfall and less predictive start and end of the rainy season) that emerges from our empirical study in rice does not seem to be covered by those models and calls for further model development. Implication of our finding for the development of crop varieties resilient to climate changes is the application of principles enunciated as early as the 16th century by Olivier de Serres (1600) “some level of intra-varietal diversity is guarantee for stability.” Such stability requires moving from the conventional paradigm of creation of uniform and genetically stable cultivars toward evolutionary plant breeding (reviewed in Döring et al., 2011), i.e. crop populations with some level of genetic diversity capable of adapting to the conditions under which they are grown.

## Data availability

- Passport and phenotypic data of the rice accessions studied are provided supplementary table 1.
- Genotypic data can be downloaded in HapMap format from http://tropgenedb.cirad.fr/tropgene/JSP/interface.jsp?module=RICE study Genotypes, study type G-panel_GBS_data.

## Authors’ contributions

NA & MBB conceived the study and assembled the rice accessions; JF produced the genotypic data. MAT produced the phenotypic data; MN Contributed to genome scan analysis; NA analysed the data and wrote the manuscript.

## Benefit-Sharing Statement

This work was undertaken in the framework of collaboration between scientists from IRAG-Guinea and Cirad-France. All collaborators are included as co-authors. The results of research have been shared with the provider communities though a number of meeting with farmers community and extension services. The aim of the present paper is to share the results with the broader scientific community. The rice accessions collected were sent to the Africa-Rice’s gene-back for long-term conservation. In relation with this research, an IRAG young scientist (Mamadou Aminata TOURE, co-author of the present manuscript) spent one year in France and for his Mater degree in genetics & plant breeding.

## Supplementary material

**S Table 1**: Passport data of the 530 rice accessions from the collected campaigns of 1979-82 and 2011, genotyped for the present study.

**S Table 2**: Variation of days to heading (DTHD) phenotyped in 2012 (Year-1) and 2013 (Year-2).

**S Table 3**: Molecular diversity and population parameters.

**S Table 4**: Variability of decay of pairwise linkage disequilibrium with distance between markers among the 12 chromosomes in five rice populations.

**S Table 5**: Loci under selection detected by heterozygosity-base method (Excoffier et al. 2009) and by drift-based method (Navascués et al. 2020), in *O. glaberrima* (*Og*) and *O. sativa japonica* groups.

**S Table 6**: Detailed results of gene enrichment analyses using Agri-Go (Tian et al. 2017; http://bioinfo.cau.edu.cn/agriGO/) and Panther-GO (Mi et al. 2019; http://GeneOntology.org) enrichment analysis tools.

**S Table 7**: Presence of QTL and genes involved in days to heading (DTHD), on 31 chromosomic segments bearing the 31 independent loci under selection.

**S Figure 1**: Area and road map of the two collect campaigns of rice samples in Guinea. Adapted from Bezançon et al. (1983).

**S Figure 2**: Pattern of deviation of the annual rainfall total (ART) and annual rainfall total during crop growing season (GSART) from the normal reference, during the 1961-2010 period, in Kankan (10°23’01.65”N, 9°18’18.72”W) and N’zérekoré (7°48’53.2”N, 8°42’14.11”W) sites of Guinea.

**S Figure 3**: Unweighted neighbor-joining tree of simple matching distances constructed from genotypes at 1,130 SNP loci, for *O. glaberrima* (*Og*) *O. sativa indica* (*Osi*) and *O. sativa japonica* (*Osj*) groups. Accessions from the first collect time (*Og*-1, *Osi*-1 and *Osj*-1) are shown in black.

**S Figure 4**: Linkage disequilibrium between SNP loci under selection in *O. glaberrima* and *O. sativa japonica* (74 and 110 SNP loci respectively). Triangle above and below the bisectrix represent the r^2^ and the r^2^ p-value respectively.

## References

Aguirre-Liguori JA, Ramírez-Barahona S, Gaut BS (2021) The evolutionary genomics of species’ responses to climate change. Nat Ecol Evol 5: 350–1360. https://doi.org/10.1038/s41559-021-01526-9

Anderson JT, Willis JH, Mitchell-Olds T (2011) Evolutionary genetics of plant adaptation. Trends Genet. 27(7): 258–266. http://dx.doi.org/10.1016/j.tig.2011.04.001

Bailey SF, Bataillon T (2016) Can the experimental evolution programme help us elucidate the genetic basis of adaptation in nature? Molecular Ecology 25: 203–218.

Barry MB, Diagne A, Sogbossi MJ, Pham JL, Diawara S, Ahmadi N (2009). Recent changes in varietal diversity of rice in Guinea. Plant genetic resources: Characterization and Utilization, 7 (1): 63–71.

Dang X, Yang Y, Zhang Y, Chen X, Fan Z, et al. (2020) OsSYL2AA, an allele identified by gene-based association, increases style length in rice (Oryza sativa L.). The Plant Journal 104: 1491–1503. http://dx.doi.org/10.1111/tpj.15013

Barry MB, Pham JL, Courtois B, Billo C, Ahmadi N (2007) Rice genetic diversity at farm and village levels and genetic structure of local varieties reveal need for in situ conservation. Genetique Resources & Crop Evolution 54: 1675–1690.

Barry MB, Pham JL, Courtois B, Billo C, Ahmadi N (2007c) Rice genetic diversity at farm and village levels and genetic structure of local varieties reveal need for in situ conservation. Genetic Resources & Crop Evolution 54: 1675–1690

Barry MB, Pham JL, Noyer JL, Courtois B, Billot C, Ahmadi N (2007b). Implications for in situ genetic resource conservation from the eco-geographical distribution of rice genetic diversity in Maritime Guinea. Plant Genetic Resources: Characterization and Utilization 5(1): 45–54.

Beaumont MA, Nichols RA (1996) Evaluating loci for use in the genetic analysis of population structure. Proceedings of the Royal Society London B 263: 1619–1626.

Bezançon G, de Kochko A, Koffi Goli (1983) Cultivated and wild species rice collected in Guinea. Plant Gent Res Newslett 57: 43–46

Bradbury PJ, Zhang Z, Kroon DE, Casstevens TM, Ramdoss Y, Buckler ES (2007) TASSEL: software for association mapping of complex traits in diverse samples. Bioinformatics 23(19): 2633–2635. https://doi.org/10.1093/bioinformatics/btm308

Cortés AJ, López-Hernández F (2021) Harnessing CropWild Diversity for Climate Change Adaptation. Genes, 12: 783. https://doi.org/10.3390/

Cruzan MB, Hendrickson EC (2020) Landscape Genetics of Plants: Challenges and Opportunities. Plant Comm. 1: 100100. https://doi.org/10.1016/j.xplc.2020.100100

Davis KF, Gephart JA, Emery KA, Leach AM, Galloway JN, D’Odorico P. (2016) Meeting future food demand with current agricultural resources. Glob. Environ. Chang. 39: 125–132.

De Navascués M, Becheler A, Gay L, Ronfort J, Loridon K, Vitalis R (2021) Power and limits of selection genome scans on temporal data from a selfing population. Peer Community Journal, V.1:e37. https://peercommunityjournal.org/articles/10.24072/pcjournal.47/

De Navascués M, Vitalis R (2020) DriftTest v1.0.5. a computer program to detect selection from temporal genetic differentiation in partially selfing populations. Zenodo. https://doi.org/10.5281/zenodo.4034846

Excoffier L, Lische HEI (2010) Arlequin suite ver 3.5: a new series of programs to perform population genetifcs analyses under Linux and Windows. Mol Ecology Ressources 10: 564–567. http://dx.doi.org/10.1111/j.1755-0998

Excoffier L, Hofer T, Foll M (2009) Detecting loci under selection in a hierarchically structured population. Heredity: 103: 285–298. http://doi.org/10.1038/hdy.2009.74

Fahad S, Bajwa AA, Nazir U, Anjum SA, Farooq A, et al. (2017) Crop Production under Drought and Heat Stress: Plant Responses and Management Options. Front. Plant Sci. 8:1147. http://doi.org/10.3389/fpls.2017.01147

Fang M, Zhou Z, Zhou X, Yang H, Li M, Li H (2019) Overexpression of OsFTL10 induces early flowering and improves drought tolerance in Oryza sativa L. PeerJ 7:e6422 http://doi.org/10.7717/peerj.6422

Footitt S, Hambidge AJ, Finch-Savage WE (2021) Changes in phenological events in response to a global warming scenario reveal greater adaptability of winter annual compared with summer annual arabidopsis ecotypes. Annals of Botany 127: 111–122. http://doi.org/10.1093/aob/mcaa141

Franks SJ, Hoffmann AA (2012) Genetics of climate change adaptation. Annual Review of Genetics, 46: 185–208.

Franks SJ, Weber JJ, Aitken SN (2014) Evolutionary and plastic responses to climate change in terrestrial plant populations. Evol Appl. 7: 123–139. https://doi.org/10.1111/eva.12112

Franks SJ, Weis AE (2008) A change in climate causes rapid evolution of multiple life-history traits and their interactions in an annual plant. J. Evol. Biol. 21: 1321–34

Gao H, Jin MN, Zheng XM, Chen J, Yuan DY, et al. (2014) Days to heading 7, a major quantitative locus determining photoperiod sensitivity and regional adaptation in rice. Proc Natl Acad Sci USA 111: 16337– 16342. https://doi.org/10.1073/pnas.1418204111

Gay L, Dhinaut J, Jullien M, Vitalis R, Navascués M, Ranwez V, Ronfort J (2021) Evolution of flowering time in a selfing annual plant: Roles of adaptation and genetic drift. bioRxiv,2020.08.21.261230, ver. 4 recommended and peer-reviewed by Peer Community in Evolutionary Biology. https://doi.org/10.1101/2020.08.21.261230

Goldringer I, Bataillon T (2004) On the distribution of temporal variations in allele frequency consequences for the estimation of effective population size and the detection of loci undergoing selection. Genetics 168: 563–568. http://doi.org/10.1534/genetics.103.025908

Gray SB, Brady SM (2016) Plant developmental responses to climate change. Developmental Biology 419: 64–77. http://doi.org/10.1016/j.ydbio.2016.07.023

Guo T, Mu Q, Wang J, Vanous AE, Onogi A, Iwata H, Li X, Yu J (2020) Dynamic effects of interacting genes underlying rice flowering-time phenotypic plasticity and global adaptation. Genome Research 30: 673– 683. http://doi.org/10.1101/gr.255703.119

Hansen MM, I Olivieri, DM Waller, EE Nielsen, The Gem working group (2012). Monitoring adaptive genetic responses to environmental change. Molecular Ecology 21: 1311–1329.

Hansen MM, Olivieri I, Waller DM, Nielsen EE, the GeM working group (2012) Monitoring adaptive genetic responses to environmental change. Molecular Ecology http://doi.org/10.1111/j.1365-294X.2011.05463.x

Hartfield M, Bataillon T, Glémin S (2017) The evolutionary interplay between adaptation and self-fertilization. Trends in Genetics, 33(6): 420–431. http://dx.doi.org/10.1016/j.tig.2017.04.002

Haussmann BIG, Rattunde HF,Weltzien-Rattunde E, Traoré PSC, vom Brocke K, Parzies HK (2012) Breeding Strategies for Adaptation of Pearl Millet and Sorghum to Climate Variability and Change in West Africa. J. Agonomy & Crop science 198(5): 327–339. https://doi.org/10.1111/j.1439-037X.2012.00526.x

Hermisson J, Pennings PS (2017) Soft sweeps and beyond: understanding the patterns and probabilities of selection footprints under rapid adaptation. Methods in Ecology and Evolution, 8: 700–716.

Hirose F, Shinomura T, Tanabata T, Shimada H, Takano M (2006) Involvement of Rice Cryptochromes in De-etiolation Responses and Flowering. Plant Cell Physiol. 47(7): 915–925. http://doi.org/10.1093/pcp/pcj064

Hoffman AA, Willi Y (2008). Detecting genetic responses to environmental change. Nature Reviews, Genetics, 9: 421–432.

Hoffmann AA, Sgrò CM (2011) Climate change and evolutionary adaptation. Nature, 470: 479–485. https://doi.org/10.1038/nature09670

Hu Y, Zhou X, Zhang B, Li S, Fan X, Zhao H, Zhang J, Liu H, He Q, Li Q, Ayaad M, You A, Xing Y(2021) OsPRR37 Alternatively Promotes Heading Date Through Suppressing the Expression of Ghd7 in the Jaonica Variety Zhonghua 11 under Natural Long-Day. Rice 14: 20. https://doi.org/10.1186/s12284-021-00464-1

Ishikawa R, Aoki M, Kurotani K, Yokoi S, Shinomura T, Takano M, Shimamoto K (2011) Phytochrome B regulates Heading date 1 (Hd1)-mediated expression of rice florigen Hd3a and critical day length in rice. Mol Genet Genom 285: 461–470. https://doi.org/10.1007/s00438-011-0621-4

Izawa T, Mihara M, Suzuki Y, Gupta M, Itoh H, et al. (2011) Os-GIGANTEA confers robust diurnal rhythms on the global transcriptome of rice in the field. Plant Cell 23: 1741–1755. https://doi.org/10.1105/tpc.111.083238

Kelly M (2019) Adaptation to climate change through genetic accommodation and assimilation of plastic phenotypes. Phil. Trans. R. Soc. B 374: 20180176. http://dx.doi.org/10.1098/rstb.2018.0176

Lamb P J (1982) “Persistence of Sub-Saharan Drought.” Nature 299: 46–8.

Lande R (1977). Statistical tests for natural selection on quantitative characters. Evolution 31: 442–444. http://doi.org/10.2307/2407764

Liu J, Sun X, Liao W, Zhang J, Liang J, Xu W (2019) Involvement of OsGF14b Adaptation in the Drought Resistance of Rice Plants Rice 12: 82 https://doi.org/10.1186/s12284-019-0346-2

Liu X, Zhou C, Zhao Y, Zhou S, Wang W, Zhou DX (2014) The rice enhancer of zeste [E(z)] genes SDG711 and SDG718 are respectively involved in long day and short day signaling to mediate the accurate photoperiod control of flowering time. Front Plant Sci 5: 591. https://doi.org/10.3389/fpls.2014.00591

Loarie SR, Duffy PB, Hamilton H, Asner GP, Field CB, Ackerly DD (2009). The velocity of climate change. Nature, 462: 1052–1055. https://doi.org/10.1038/nature08649

Lohani N, Singh MB, Bhalla PL (2020) High temperature susceptibility of sexual reproduction in crop plants. Journal of Experimental Botany, 71(2): 555–568. http://doi.org/10.1093/jxb/erz426

Loua RT, Beavogui M, Bencherif H, Barry HB, Bamba Z, Mazodier CA (2017) Climatology of Guinea: Study of Climate Variability in N’zerekore. Journal of Agricultural Science and Technology A 7: 215–233. http://doi.org/10.17265/2161-6256/2017.04.001

Luo M, Plattena D, Chaudhury A, Peacocka WJ, Dennisa ES (2009) Expression, Imprinting, and Evolution of Rice Homologs of the Polycomb Group Genes. Molecular Plant 2: 711–723.

Messer PW, Petrov DA (2013) Population genomics of rapid adaptation by soft selective sweeps. Trends Ecol. Evol. 28: 659–669.

Mi H, Muruganujan A, Ebert D, Huang X, Thomas PD (2019) PANTHER version 14: more genomes, a new PANTHER GO-slim and improvements in enrichment analysis tools. Nucleic Acids Research, 47: 419–426. http://doi.org/10.1093/nar/gky103

Miller-Rushing AJ, Primack RB (2008) Global warming and flowering times in Thoreau’s Concord: a community perspective. Ecology 89:332–41

Mouradov A, Cremer F, Coupland G (2002) Control of Flowering Time: Interacting Pathways as a Basis for Diversity. The Plant Cell, S111–S130.

Murakami M, Matsushika A, Ashikari M, Yamashino T, Mizuno T (2005) Circadian-aAssociated Rice Pseudo Response Regulators (osprrs): Insight into the Control of Flowering Time. Biosci. Biotechnol. Biochem. 69 (2): 410–414.

Oka H, Morishima H (1967) Variations in the breeding systems of a wild rice, oryza perennis. Evolution 21: 249–258. https://doi.org/10.1111/j.1558-5646.1967.tb00153

Orr HA (2002) The population genetics of adaptation: the adaptation of DNA sequences; Evolution 56(7): 1317–1330.

Orr HA (2005) The genetic theory of adaptation: a brief history. Nature reviews genetics 6: 119–127.

Pauls SU, Nowak C, Bálint M, Pfenninger M (2013) The impact of global climate change on genetic diversity within populations and species. Mol Ecol. 22: 925–946. https://doi.org/10.1111/mec.12152

Perrier X, Jacquemoud-Collet JP (2006) DARwin software (http://darwin.cirad.fr/darwin)

Pironon S, Etherington TR, Borrell JS, Kühn N, Macias-Fauria M, et al. (2019). Potential adaptive strategies for 29 sub-Saharan crops under future climate change. Nat. Clim. Chang. 9: 758–763. https://doi.org/10.1038/s41558-019-0585-7

Prevéy JS (2020) Climate change: flowering time may be shifting in surprising ways. Current Biology 30: 112–133. https://doi.org/10.1016/j.cub.2019.12.009

Purwestri Y A, Ogaki Y, Tamaki S, Tsuji H, Shimamoto K (2009) The 14-3-3 protein GF14c acts as a negative regulator of flowering in rice by interacting with the florigen Hd3a. Plant Cell Physiol. 50: 429– 438. http://doi.org/10.1093/pcp/pcp012

Quiroz S, Yustis JC, Chávez-Hernández EC, Martínez T, Sanchez M, Garay-Arroyo A, Álvarez-Buylla ER, García-Ponce B (2021) Beyond the Genetic Pathways, Flowering Regulation Complexity in Arabidopsis thaliana. Int. J. Mol. Sci. 22: 5716. https://doi.org/10.3390/ijms22115716

Radanielina T, Ramanantsoanirina A, Raboin L-M, Ahmadi N (2013) Déterminants de la diversité variétale du riz dans la région de Vakinankaratra, Madagascar. Cah Agric 22(5): 442–449. https://doi.org/10.1684/agr.2013.0648

Reush and Wood (2007) Molecular ecology of global change. Molecular ecology, 16:3973–3992.

Rojas M, Lambert F, Ramirez-Villegas J, Challinor A J (2019). Emergence of robust precipitation changes across crop production areas in the 21st century. Proc. Natl Acad. Sci. USA, 116: 6673–6678

Sahadevan PC, Namboodiri KMN (1963). Natural crossing in rice. Proc. Indian Acad. Sci. Societ., 58(2): 176–185.

Snowdon RJ, Wittkop B, Chen T-W, Stahl A (2021) Crop adaptation to climate change as a consequence of long?term breeding. TAG 134: 1613–1623. https://doi.org/10.1007/s00122-020-03729-3

Springate DA, Scarcelli N, Rowntree J, Kover PX (2011) Correlated response in plasticity to selection for early flowering in Arabidopsis thaliana. J Evol Biol. 24: 2280–2288. https://doi.org/10.1111/j.1420-9101.2011.02360.x

Sultan B, Defrance D, Iizumi T (2019) Evidence of crop production losses in West Africa due to historical global warming in two crop models. Sci. Rep. 9: 1–15. https://doi.org/10.1038/s41598-019-49167-0

Takano M, Inagaki N, Xie X, Yuzurihara N, Hihara F, et al. (2005) Distinct and cooperative functions of phytochromes A, B, and C in the control of deetiolation and flowering in rice. Plant Cell 17: 3311–3325. https://doi.org/10.1105/tpc.105.035899

Tian T, Liu Y, Yan H, You Q, Yi X, Du Z, Xu W, Su Z (2017) agriGO v2.0: a GO analysis toolkit for the agricultural community, 2017 update. Nucleic Acids Research, 45 W122–W129. https://doi.org/10.1093/nar/gkx382

Wanga J, Hub J, Qianb Q, Xuea HW (2013) LC2 and OsVIL2 Promote Rice Flowering by Photoperoid-Induced Epigenetic Silencing of OsLF. Molecular Plant 6(2): 514–527.

Wassmann R, Jagadish SVK, Peng SB, Sumfleth K, Hosen Y, Sander BO (2010) Rice production and global climate change: scope for adaptation and mitigation activities. Wassmann R, editor. Proceedings of the Workshop Advanced Technologies of Rice Production for Coping with Climate Change: ‘No Regret’ Options for Adaptation and Mitigation and their Potential Uptake held on 23-25 June 2010 in Los Baños, Philippines. IRRI Limited Proceedings No. 16. Los Baños (Philippines): International Rice Research Institute. 81 p.

Watson D (2019) Adaption to Climate Change: Climate Adaptive Breeding of Maize, Wheat and Rice. In: Sarkar A, Sensarma S, vanLoon G (eds) Sustainable Solutions for Food Security. Springer, Cham. https://doi.org/10.1007/978-3-319-77878-5_4

Wei H, Wang X, Xu H, Wang L (2020) Molecular basis of heading date control in rice. aBIOTECH 1: 219– 232. https://doi.org/10.1007/s42994-020-00019-w

Weir BS, Cockerham CC (1984) Estimating F-statistics for the analysis of population structure. Evolution 38: 1358–1370.

Wellmer F, Riechmann JL (2010) Gene networks controlling the initiation of flower development. Trends in Genetics, 26: 519–527.

WMO/OMM (1996) Climatological normal (CLINO) for the period 1961-1990. WMO Series n°847. SBN 10: 9263008477

Wright S (1965) The interpretation of population structure by F-statistics with special regard to systems of mating. Evol 19: 395–420.

Yu J, Jiang M, Guo C (2019) Crop Pollen Development under Drought: From the Phenotype to the Mechanism. Int. J. Mol. Sci. 20, 1550. https://doi.org/10.3390/ijms20071550

Zhang S, Jin Y, Hao H, Liang S, Ma X, Luan W(2020) Characterization and identification of OsFTL8 gene in rice; Plant Biotechnology Reports 14: 683–694. https://doi.org/10.1007/s11816-020-00644-3

