## Supplementary figures and images for "Genome scan of landrace populations of the self-fertilizing crop species rice, collected across time, revealed climate changes’ selective footprints in the genes network regulating flowering time"

### S Figure 1.tif

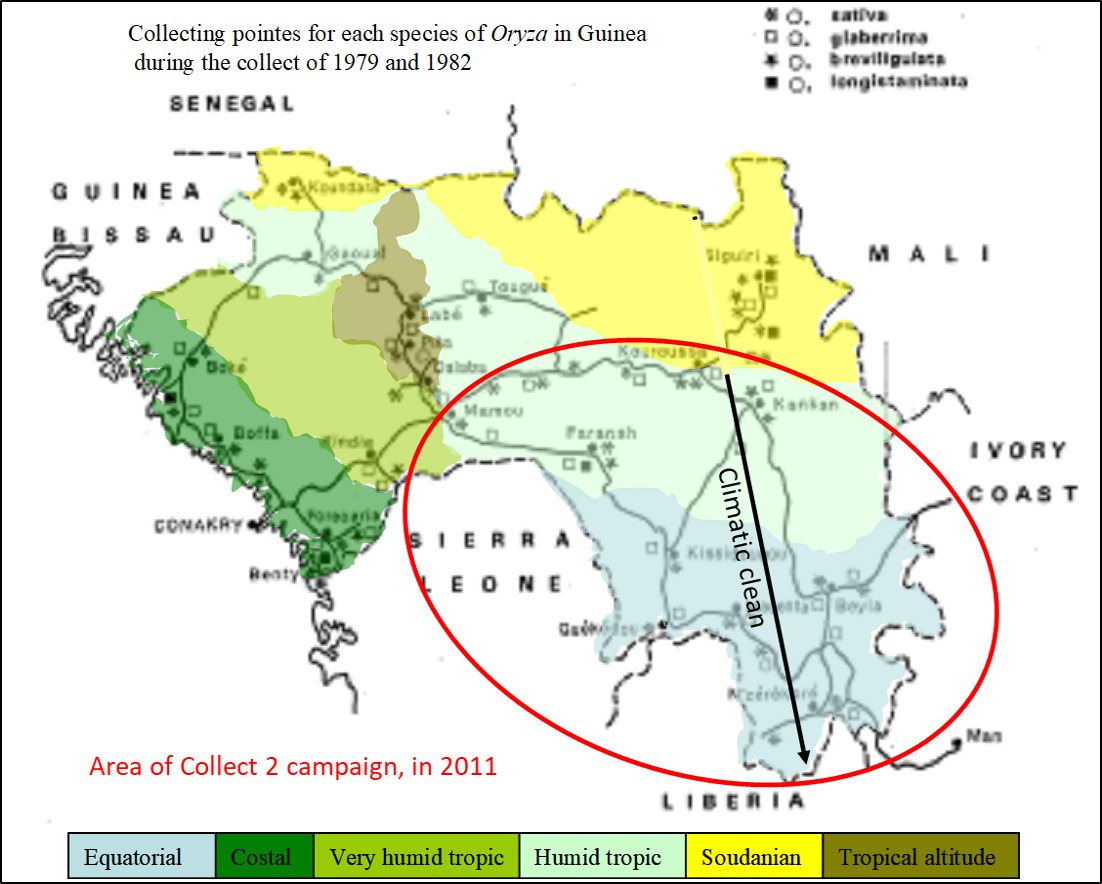

### S Figure 2.tif

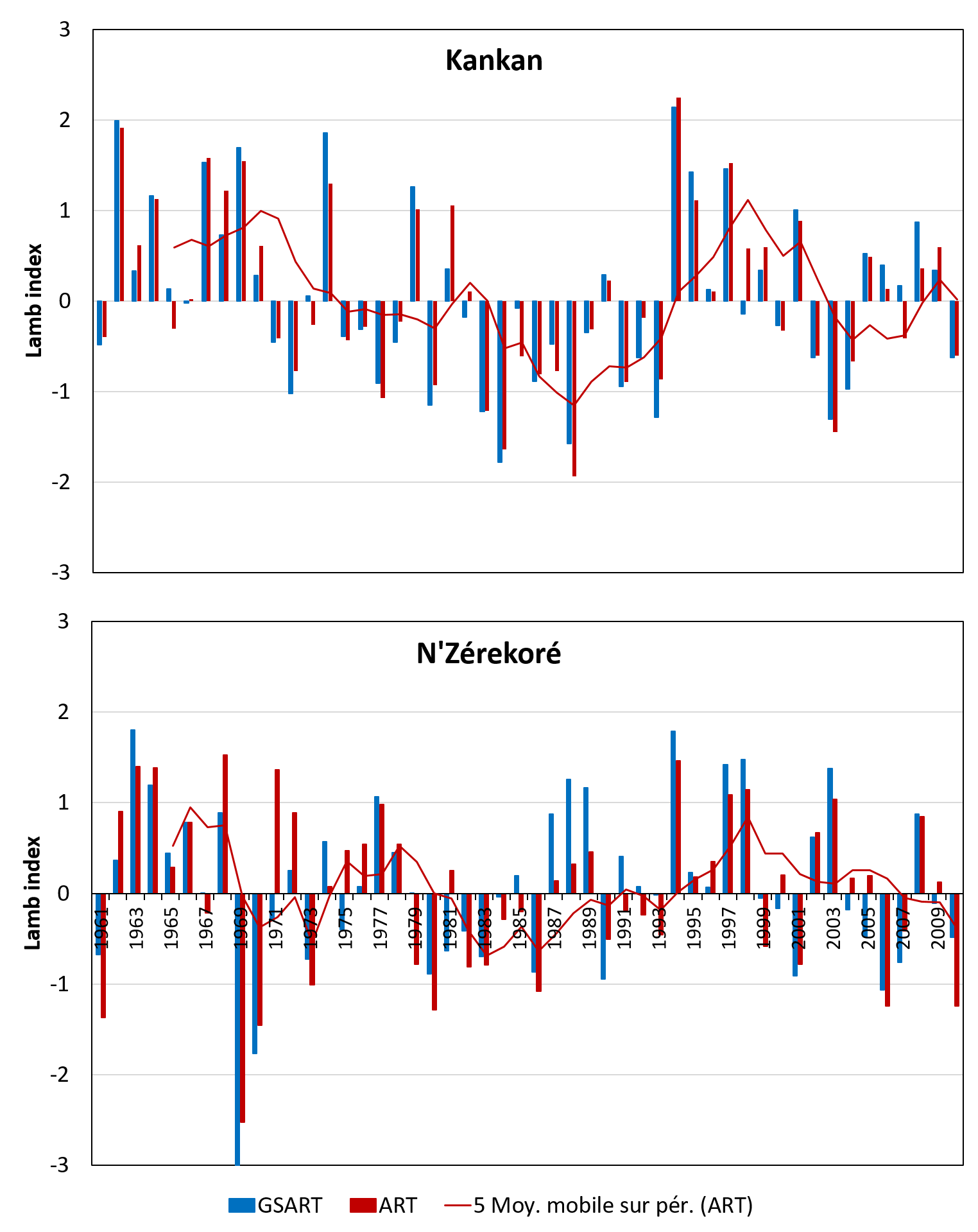

### S Figure 3.tif

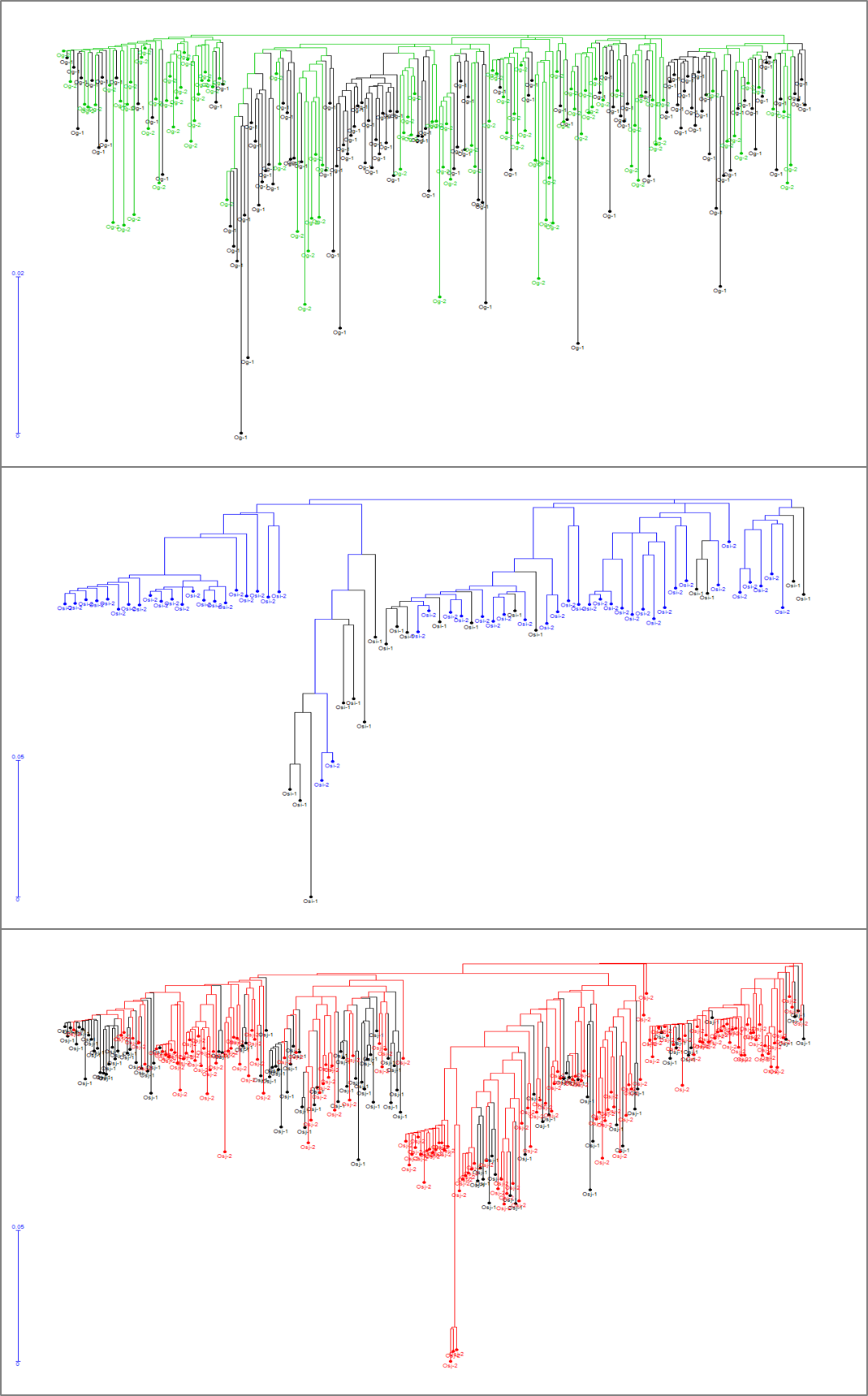

### S Figure 4.tif

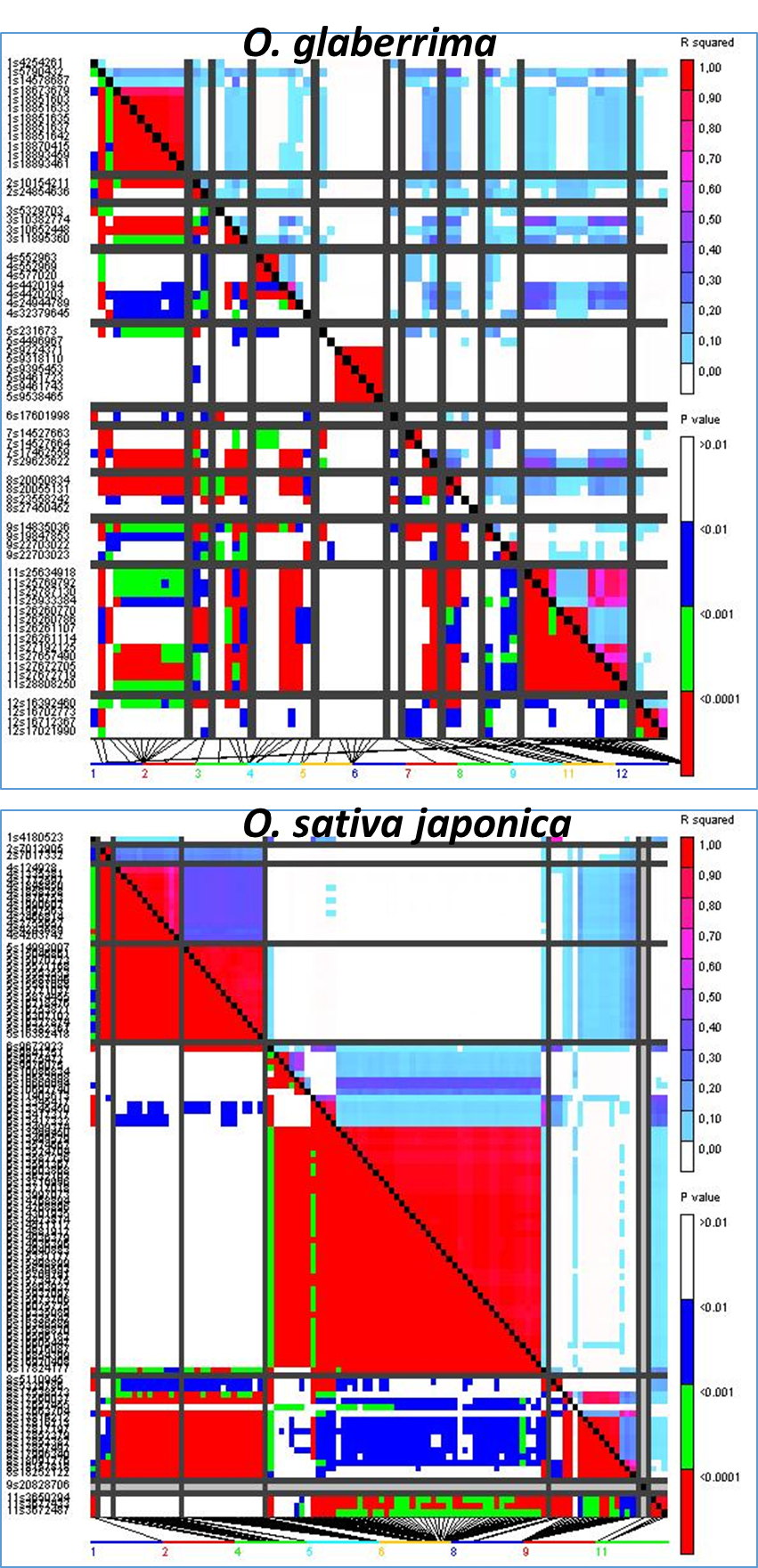
